# Discovery of diverse human BH3-only and non-native peptide binders of pro-apoptotic BAK indicate that activators and inhibitors use a similar binding mode and are not distinguished by binding affinity or kinetics

**DOI:** 10.1101/2022.05.07.491048

**Authors:** Fiona Aguilar, Stacey Yu, Robert A. Grant, Sebastian Swanson, Dia Ghose, Bonnie G. Su, Kristopher A. Sarosiek, Amy E. Keating

## Abstract

Apoptosis is a programmed form of cell death important for the development and maintenance of tissue homeostasis. The BCL-2 protein family controls key steps in apoptosis, dysregulation of which can lead to a wide range of human diseases. BCL-2 proteins comprise three groups: anti-apoptotic proteins, pro-apoptotic proteins, and BH3-only proteins. BAK is one of two pro-apoptotic proteins, and previous work has shown that binding of certain BH3-only proteins such as truncated BID (tBID), BIM, or PUMA to BAK leads to mitochondrial outer membrane permeabilization, the release of cytochrome c, and ultimately cell death. This process, referred to as *activation*, involves the BH3-stimulated conversion of BAK from monomer to dimer and then to oligomers that promote membrane disruption. Crystal structures of putative intermediates in this pathway, crosslinking data, and *in vitro* functional tests have provided insights into the activation event, yet the sequence-function relationships that make some but not all BH3-only proteins function as activators remain largely unexamined. In this work, we used computational protein design, yeast surface-display screening of candidate BH3-like peptides, and structure-based energy scoring to identify ten new binders of BAK that span a large sequence space. Among the new binders are two peptides from human proteins BNIP5 and PXT1 that promote BAK activation in liposome assays and induce cytochrome-c release from mitochondria, expanding current views of how BAK-mediated cell death may be triggered in cells. High-resolution crystal structures and binding experiments revealed a high degree of similarity in binding geometry, affinity, and association kinetics between peptide activators and inhibitors, including peptides described previously and those identified in this work. We propose a model for BAK activation that is based on differential engagement of BAK monomers vs. the BAK activation transition state that integrates our observations with previous reports of BAK binders, activators, and inhibitors.

## INTRODUCTION

BAK is a pro-apoptotic member of the BCL-2 protein family (Chittenden et al., 1995). It plays a key role in regulating apoptosis, a programmed form of cell death that is important for development and tissue homeostasis, and its dysregulation can lead to a range of diseases (Duque-Parra, 2005) (Kerr et al., 1972) (Lindsten et al., 2000) (R. Singh et al., 2019). For example, BAK is implicated in hematopoietic regulation through the clearance of mature T and B lymphocytes (Takeuchi et al., 2005) (Rathmell et al., 2002). Moreover, over-expression of BAK has been found in human brains from Alzheimer’s patients (Kitamura et al., 1998). Age- and tissue-specific differences in expression levels of BAK and the related protein BAX give rise to differences in sensitivity to cancer therapeutics (Sarosiek et al., 2017).

In an irreversible step toward cell death, BAK and BAX promote apoptosis via mitochondrial outer membrane permeabilization (MOMP) in response to a variety of signals (Tait & Green, 2010). Structural, biochemical, and cell biological studies have illuminated some aspects of the mechanism of membrane permeabilization by BAK and BAX, yet a full understanding remains elusive due to the dynamic nature of these proteins, the lipid environment in which they function, and the complex multilayered BCL-2 network that provides regulatory control (Sandow et al., 2021) (Delbridge et al., 2016). In this study, we addressed early steps in the activation of BAK in which the binding of certain BCL-2 family members to form heterodimers leads to a conformation change that is required for downstream steps including BAK dimerization and higher-order oligomerization.

The BCL-2 family can be subdivided into three groups: anti-apoptotic proteins, pro-apoptotic proteins, and BH3-only proteins. All family members contain a <u>B</u>CL-2 <u>H</u>omology <u>3</u> (BH3) motif (Aouacheria et al., 2013); the anti-apoptotic and pro-apoptotic proteins additionally contain other conserved motifs (BH1, BH2, BH4) (Youle & Strasser, 2008). Anti-apoptotic and pro-apoptotic proteins include a C-terminal transmembrane segment that can anchor them in the <u>m</u>itochondrial <u>o</u>uter <u>m</u>embrane (MOM) and 8 helices that adopt a globular fold localized to the cytoplasmic face of the membrane (Youle & Strasser, 2008). This globular domain contains a hydrophobic groove that can be bound in trans by the BH3 motif regions of other MOM-anchored BCL-2 proteins (Sattler et al., 1997). The pro-apoptotic proteins include BAK, BAX, and less-studied BOK (Moldoveanu & Czabotar, 2019). BAK and BAX are structurally and functionally very similar, although differences in the mechanistic details of their activation and function have been reported (Sarosiek et al., 2013) (Czabotar et al., 2013) (Brouwer et al., 2014) (Garner et al., 2016).

BH3-only proteins can be classified based on their functions as either ‘sensitizers’ or ‘activators’. All known BH3-only proteins bind to anti-apoptotic BLC-2 proteins with high affinity (K_d_ ≤ 10 nM), competitively inhibiting their engagement of BAK and BAX, and thus *sensitizing* the cell to undergo apoptosis (Willis et al., 2005) (Willis et al., 2007) (Zheng et al., 2016). Only three human proteins contain BH3 motifs that are known to bind directly to BAK and induce a conformational change that leads to BAK dimerization and then MOMP: truncated BID (tBID), BIM, and PUMA; we refer to these as ‘activators’ (Kim et al., 2009) (Llambi et al., 2011) (Dai et al., 2014). The sequence and structural features that distinguish a sensitizer from an activator are unknown, as is the mechanism of BH3-binding induced activation.

There is controversy in the field regarding the requirement of direct activation for BAK-dependent cell death. There is clear evidence from biochemical studies using liposomes that BH3 peptides can directly activate both BAK and BAX (Brouwer et al., 2017) (Gavathiotis et al., 2008) (G. Singh et al., 2022). Adding such peptides to mitochondria or permeabilized cells with intact mitochondria also induces MOMP (Sarosiek et al., 2013) (Moldoveanu et al., 2013). The indirect activation model, on the other hand, posits that the binding of BH3 proteins to anti-apoptotic proteins localized at the mitochondria is sufficient to allow unimpeded, spontaneous activation of BAK or BAX, perhaps triggered by the mitochondrial membrane itself (Huang et al., 2019). In this model, activator BH3-only proteins are not required for binding to BAK or BAX. The strongest evidence for this model comes from experiments involving induced expression of sensitizer BAD (a binder of anti-apoptotic BCL-2, BCL-x_L_, and BCL-W, but not BAK or BAX) in cells lacking all other BH3-only proteins and anti-apoptotic MCL-1 (Huang et al., 2019). In this setting, expression of BAD alone is sufficient to induce BAX-mediated MOMP (Huang et al., 2019). A key question is whether all activating proteins were depleted in these cell lines, or whether unknown activators might have had some role in these experiments. Further studies looking at BAK in addition to BAX and using physiological concentrations of protein will help clarify the implications of this work.

Regardless of the relative roles of direct vs. indirect BAK activation *in vivo*, direct activation is robust and well-established in *in vitro* assays consisting of liposomes, recombinant protein, and BH3 peptides. These observations suggest that BAK can be activated by endogenous or exogenous BH3 activators in cells, regardless of whether this is required for cell death. In this work, we investigated biochemical requirements for direct activation.

Structures and crosslinking studies have provided a working model to describe some of the steps involved in BAK activation, i.e., the path that is taken from a membrane-tethered BAK monomer (PDB: 2IMS) to a BH3-bound monomer (PDB: 5VX0) and ultimately a membrane-embedded association of BAK homodimers (**Figure 1**) (Brouwer et al., 2017) (Kuwana et al., 2002) (Sandow et al., 2021). Crystal structures and crosslinking data support a conformational change in which the “latch” region (BAK α6-α8) disengages from the BAK “core” region (α2-α5) (PDB: 5VWV) (Brouwer et al., 2014). Two activated BAK monomers can then exchange BH3 helices and form stable BH3:groove homodimers (PDB:7K02) that then associate with other dimers and ultimately give rise to MOMP (Brouwer et al., 2014) (Birkinshaw et al., 2021).

**Figure 1.**
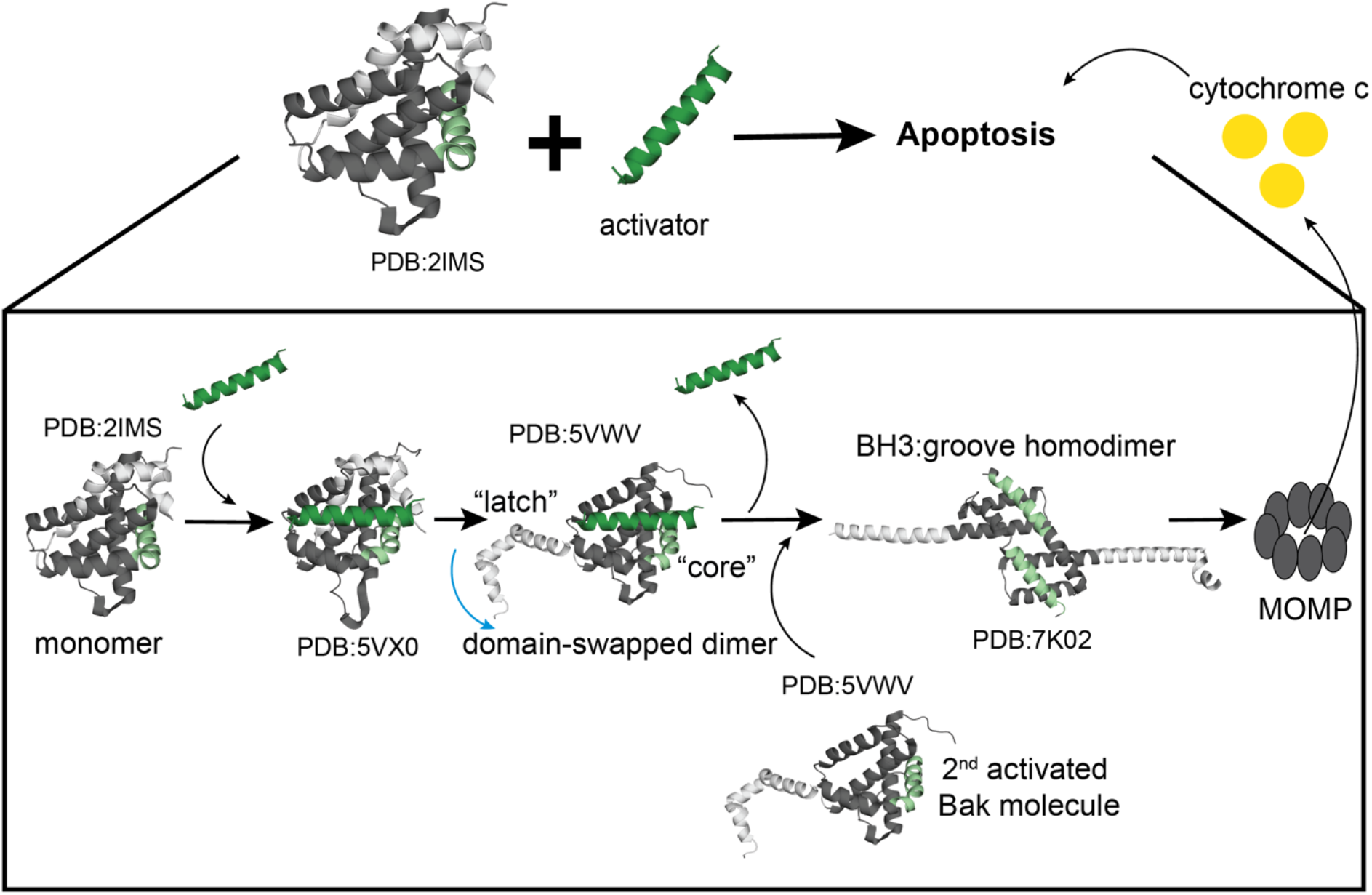
Model of BAK activation based on previously published putative intermediate crystal structures. The current model for BAK activation occurring on the mitochondrial outer membrane involves the binding of a BH3 segment (dark green) from an activating protein to monomeric BAK (grey) (PDB:2IMS), as observed in the structure of BAK bound to a modified BIM BH3 peptide (PDB:5VX0). Binding leads to the release of the BAK “latch” (α6-α8) (light grey) from the “core” (α2-α5) (dark grey) as observed in domain-swapped structures of BAK (PDB:5VWV). Subsequent conformational changes lead to a “BH3:groove homodimer” (PDB:7K02) formed by the core regions of two BAK monomers that exchange BH3 helices (light green).

Given the therapeutic potential of targeting BAK or BAX to treat cancer or neurodegenerative diseases, there is great interest in developing tools to modulate the functions of these proteins and understand their complex activation mechanisms (Reyna et al., 2017) (Pritz et al., 2017) (Niu et al., 2017)(Delft et al., 2019) (Iyer et al., 2020). For example, previous research has shown that is it possible to bind BAK and inhibit rather than activate it, using tight-binding peptides containing non-natural amino acids (Brouwer et al., 2017). Conversely, antibodies targeting the α1-α2 loop of BAK can induce MOMP, demonstrating an alternative binding site to trigger activation (Iyer et al., 2016). Other groups have developed small molecules that sensitize BAX activation (Pritz et al., 2017) as well as inhibit BAX oligomerization (Niu et al., 2017).

In this study, we explored the sequence space of peptide binders of BAK using computational structure-based design and cell-surface display screening. We obtained novel non-native BAK-binding peptides with diverse sequences. Additionally, we discovered nine previously unknown binders of BAK in the human proteome, two of which function as activators in liposome assays and mitochondrial permeabilization assays. We conducted functional, biophysical, and structural studies to dissect the differences between peptide activators and inhibitors of BAK. Our results indicate that binding affinity, binding kinetics, and binding geometry are not sufficient to clearly distinguish between the two. We speculate on the energetic requirements in each case and propose an energy-landscape model to summarize our findings and integrate them with other information in the literature.

## RESULTS

### Structure-based design and library screening identify novel peptide binders of BAK

To discover novel binders of BAK that could function as activators or inhibitors, we used computational peptide design and cell-surface screening of candidate BH3-like peptides (**Figure 2**). **D**esign with **TERM** energies (dTERMen) is a computational method that computes statistical energies from repeating structural elements in known protein structures and has been applied to design peptide binders of anti-apoptotic BCL-2 proteins (**Table S1**) (Zhou et al., 2020) (Frappier et al., 2019). Given a protein backbone structure, dTERMen mines a diverse database of known structures to identify matches to component motifs and uses the sequences of the matches to compute a Potts model that gives energy as a function of sequence. Optimizing this function rapidly provides the sequence that is predicted to be most compatible with the target structure. As input into dTERMen for BAK-binder design, we used the crystal structure of BAK in complex with Bim-h3Glg (PDB:5VX0), a peptide that contains a non-natural amino acid with a long side chain. Bim-h3Glg functions as an inhibitor of BAK activation (Brouwer et al., 2017). Our peptide designed using dTERMen, BK3, contains lysine at position 3f, which is conserved as aspartate in native BH3 peptides and is glutamate in some other BAK binders that we studied (**Figure 2A**).

**Figure 2.**
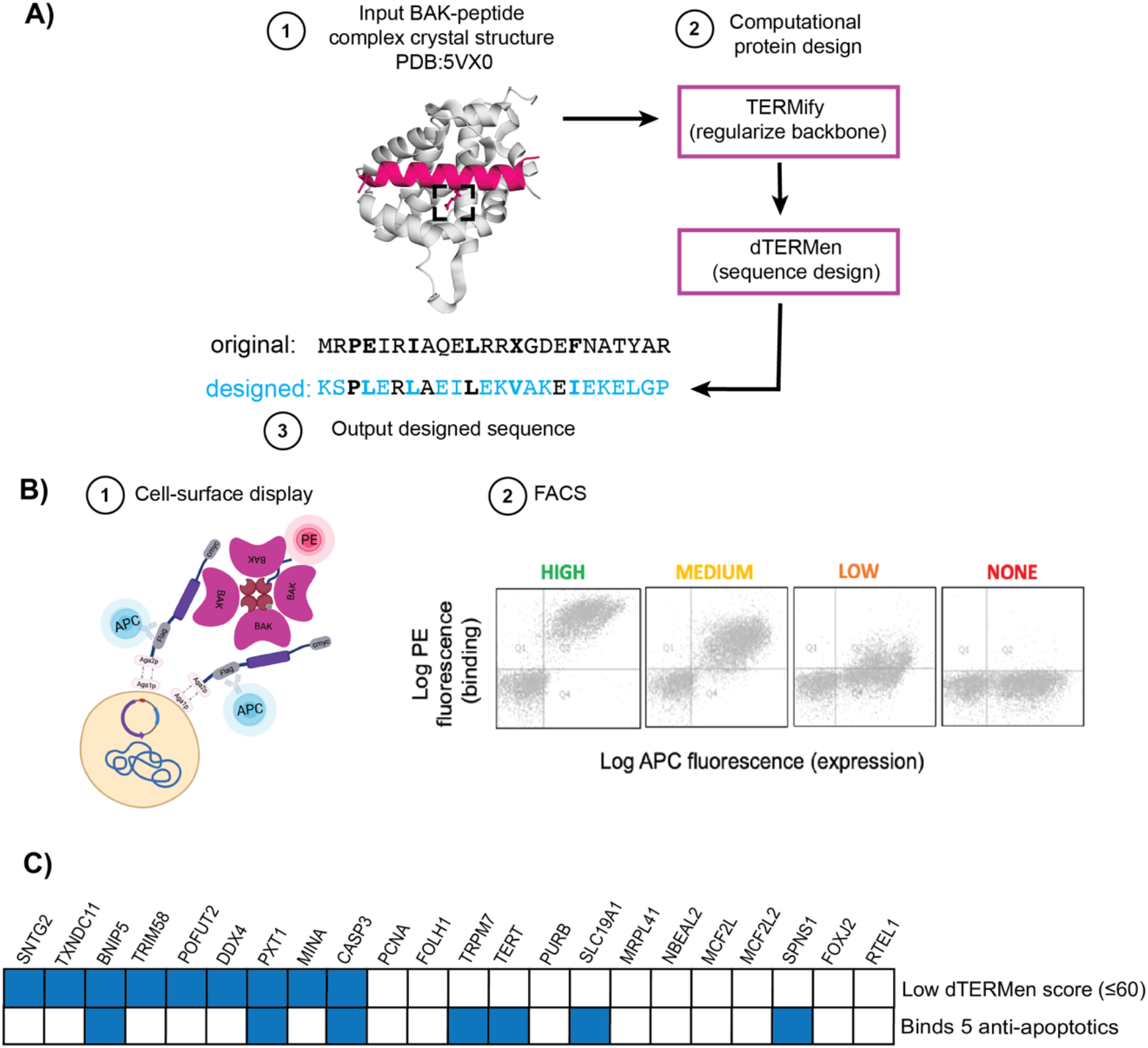
Three methods used to discover peptides binders of BAK. **A)** The structure of BAK bound to BIM-h3Glg (PDB:5VX0) was used as input for computational re-design using TERMify and dTERMen to generate a novel BAK binding sequence, BK3. A non-natural amino acid is depicted with an X in the input peptide sequence and surrounded by a dashed box in the structure. **B)** Cartoon representation of the yeast-surface display assay used to test candidate binders of BAK. Peptide expression was detected using an HA tag and binding to BAK tetramers was quantified using SAPE fluorescence. High-binding (green), medium-binding (light orange), low-binding (dark orange), and non-binding (red) peptides were determined based on binding vs. peptide expression signals using flow cytometry. Peptides were assigned manually to one of these four categories. **C)** BH3-like regions in human proteins reported by DeBartolo *et al*. were scored for binding to pro-apoptotic BAK using dTERMen. Those with favorable, low-energy scores are indicated in blue. Proteins containing peptides that bind all five of BCL-2, BCL-x_L_, MCL-1, BFL-1, and BCL-W are also indicated in blue. BH3 peptides from all of the proteins that are indicated in blue were subsequently tested for binding to BAK.

We identified additional BAK binders through yeast-surface display screening of previously published BH3-like peptide binders of anti-apoptotic BCL-2 family proteins (**Figure 2B and Table S1**). Based on the structural similarity of BAK to BCL-x_L_, MCL-1, and BFL-1, and the fact that BH3 peptides from BIM, BID, and PUMA bind to both the pro- and anti-apoptotic BCL-2 family members, we reasoned that some of these designed peptides might also bind pro-apoptotic BAK. Prior work used a monovalent yeast surface-display assay to discover binders of the anti-apoptotic proteins (Dutta et al., 2015) (Jenson et al., 2017). However, BID and BIM BH3 peptides bind to BAK much more weakly than they bind to the anti-apoptotic family members (data not shown), so we adapted the yeast-surface display framework to a multi-valent format, using pre-tetramerized BAK to enhance avidity (**Figure 2B**). We tested 28 peptides and discovered 19 that bound to tetramerized BAK on the yeast surface (**Table S1**). Binding profiles measured by Fluorescence Activated Cell Sorting (FACS) were categorized into 4 groups based on binding signal relative to peptide expression: high, medium, low, or no binding (**Figure 2B**). We performed further characterization of sequences from the high-binding group, which included dTERMen-designed peptide BK3, as described below.

### Discovery of new human BH3-only binders of BAK

In prior work, DeBartolo *et al*. scanned the human proteome to identify candidate, previously unidentified BH3-only sequences (DeBartolo et al., 2014). The survey employed structure-based scoring combined with experimental screening using SPOT arrays and identified 34 peptides that bound detectably to at least one of the five anti-apoptotic proteins MCL-1, BCL-x_L_, BCL-2, BFL-1, and BCL-W (DeBartolo et al., 2014). Given the high structural similarity between pro-apoptotic BAK and the anti-apoptotic proteins, we reasoned that some of the peptides might also bind BAK (**Figure 2C)**. Using three BAK-BH3 peptide complex structures that we solved as part of this work (see below), we aligned each of the 22 human proteome candidate BH3 sequences measured to have K_d_ ≤ 500 nM for any anti-apoptotic BCL-2 protein to the peptides in our crystal structures and calculated an energy score using dTERMen. This generated three different scores per candidate peptide (**Figure 2C and Table S2)**. We tested the binding of the top seven ranked BH3-only peptides using fluorescence anisotropy and discovered three new binders (**Table S2 and Figure S1**). Of the three, BNIP5 bound most tightly, with K_d_ = 0.4 µM. PXT1 bound with K_d_ = 11 µM, and TRIM58 was the weakest binder; with an estimated K_d_ > 20 µM (**Figure S1A**). We next expanded our test to include all of the sequences reported by DeBartolo *et al*. that bind to all 5 anti-apoptotic proteins. These included peptides from NBEAL2, SLC19A1, SPNS1, CASP3, TERT, and TRMP7 (**Figure S1B**). We discovered that all 6 sequences bound BAK weakly (K_d_ > 20 µM) as determined by fluorescence polarization binding assays (**Figure S1B**).

The binding of BH3 segments to BAK or anti-apoptotic BCL-2 family proteins requires that the BH3 region be accessible. Most BH3-containing proteins are intrinsically disordered; for BID, a cleavage event converts a folded structure that sequesters the BH3 motif into a shorter protein called tBID in which this region is exposed (McDonnell et al., 1999). AlphaFold structure predictions of known BH3-only activators BIM and PUMA show high levels of disorder, except for the BH3 region, which is predicted to be helical, as it is when it is bound to a BCL-2 anti-apoptotic protein (**Figure S2**). With the exception of CASP3, there are no solved structures for any of the human BH3-containing proteins that bind BAK. Thus, we used AlphaFold to examine predicted structures and categorized the candidates into three groups based on their per-residue pLDDT confidence score over the BH3 region, which is typically low for disordered proteins, and the predicted accessibility of the BH3 motif (**Figures S2 and S3, Tables S3 and S4)**. BNIP5 is predicted to be highly disordered, and its BH3 region is predicted to adopt an alpha helical structure with a low-confidence pLDDT score. Structure gazing indicates that the BH3 region is largely accessible and solvent exposed, as it does not form part of a tertiary structure. We reached similar conclusions for PXT1, TRIM58, BIM, and PUMA. The candidate BH3 motifs in other proteins were more structurally constrained by surrounding residues and showed less solvent accessibility, including TRPM7, TERT, and SLC19A1; these were classified as less accessible. The candidate BH3 motifs in CASP3 and SPNS1 were classified as inaccessible because major conformational changes would have to occur for a BH3-mediated interaction to occur, if the high-confidence predictions are correct.

### Novel BH3 binders of BAK function as activators or inhibitors

Liposomes loaded with recombinant proteins provide a way to study BAX and BAK activation while avoiding the influence of confounding factors such as anti-apoptotic proteins (Brouwer et al., 2017) (Gavathiotis et al., 2008) (Hockings et al., 2015). Because recombinant full-length BAK is not soluble, we used a previously published C-terminally truncated BAK construct with a His_6_ tag (BAKΔC25-His_6_) that localizes BAK to Ni^2+^-labeled liposomes (Brouwer et al., 2017). Briefly, we made 100 nm liposomes that mimicked the composition of the mitochondrial outer membrane with encapsulated fluorescent dye DPX and quencher ANTS. In this assay, dye release reports on membrane permeabilization as a proxy for MOMP. In initial experiments performed with a membrane-localized activating peptide from BID (His_6_-SUMO-BID), BAKΔC25-His_6_ but not BAK ΔN22 ΔC25 C166S (which lacks the membrane-localizing His_6_ tag) showed a response to activator (**Figure S4**). We used BAKΔC25-His_6_ in all subsequent liposome-based experiments. BAK activation involves BAK oligomerization (Cosentino et al., 2022), and we confirmed that this process is concentration-dependent: 500 nM BAKΔC25-His_6_ showed greater activation than 150 nM BAKΔC25-His_6_ (**Figure S5**).

We used this assay to test activation by a range of soluble 22-35 residue peptides (**Figure S6 and Table S5**). BH3 peptides from BAK activators BID and BIM gave concentration-dependent activation, consistent with prior reports (Brouwer et al., 2017) (Hockings et al., 2015) (G. Singh et al., 2022). Our data show that PUMA and HRK peptides give minimal activation compared to BID and BIM BH3 peptides at 2.5 µM and 5 µM. NOXA and BAD BH3 peptides did not activate up to concentrations of 5 µM.

The canonical binding grooves of pro-apoptotic proteins BAK and BAX contain 5 hydrophobic pockets referred to as h0, h1, h2, h3, and h4 (**Figure S7**). Numerous structures of BAK and BAX complexes have shown that specific positions within the BH3 peptide engage these hydrophobic pockets, and mutations to pocket-binding residues can disrupt function (Czabotar et al., 2013) (Brouwer et al., 2017). To probe the sensitivity of BAK to mutations in BH3 activators, we focused on mutations previously shown to affect the activation of BAX (**Figure S8**) (Czabotar et al., 2013). A 26-residue PUMA BH3 peptide showed weak activation at 20 µM. Substitution of native methionine 144 at position 3d with isoleucine gave greater BAK activation compared to WT. Similarly, substitution of native cysteine 25 at position 2d to isoleucine in NOXA gave greater BAK activation.

We tested our newly discovered BAK-binding peptides for BAK activation using the liposome assay (**Figure 3** and **Figure S9**). Three peptides previously designed to bind to anti-apoptotic proteins (dF8, dM2, and dM4) are very different from one another and from known activators BIM, BID, and PUMA, yet all showed an increase in fluorescence signal compared to negative controls at 1.5 h, indicating BAK activation (**Figure 3A**). A BIM BH3 peptide with solubilizing mutations (BIM-RT) was used as a positive control. dF8 and dM4 were more potent activators than BIM-RT at 1.25 µM. dM2 peptide, on the other hand, showed weaker activation compared to BIM-RT. Peptides from human proteins BNIP5 and PXT1 activated 300 nM BAKΔC25-His_6_ (**Figure 3B**). At 250 nM, BNIP5 showed greater activation than PXT1, using positive control BIM-RT at 500 nM as a reference, indicating BNIP5 is a more potent activator. To test whether our new activating peptides act synergistically or in competition with BIM BH3, we incubated 300 nM of BAKΔC25-His_6_ with 200 nM BIM-RT and increasing concentrations of dF8, dM2, or dM4 peptides. The addition of 500 nM dF8 or dM4 to 200 nM BIM-RT resulted in greater activation than observed for BIM-RT alone; this was also true but to a smaller extent for dM2 (**Figure S10**).

**Figure 3.**
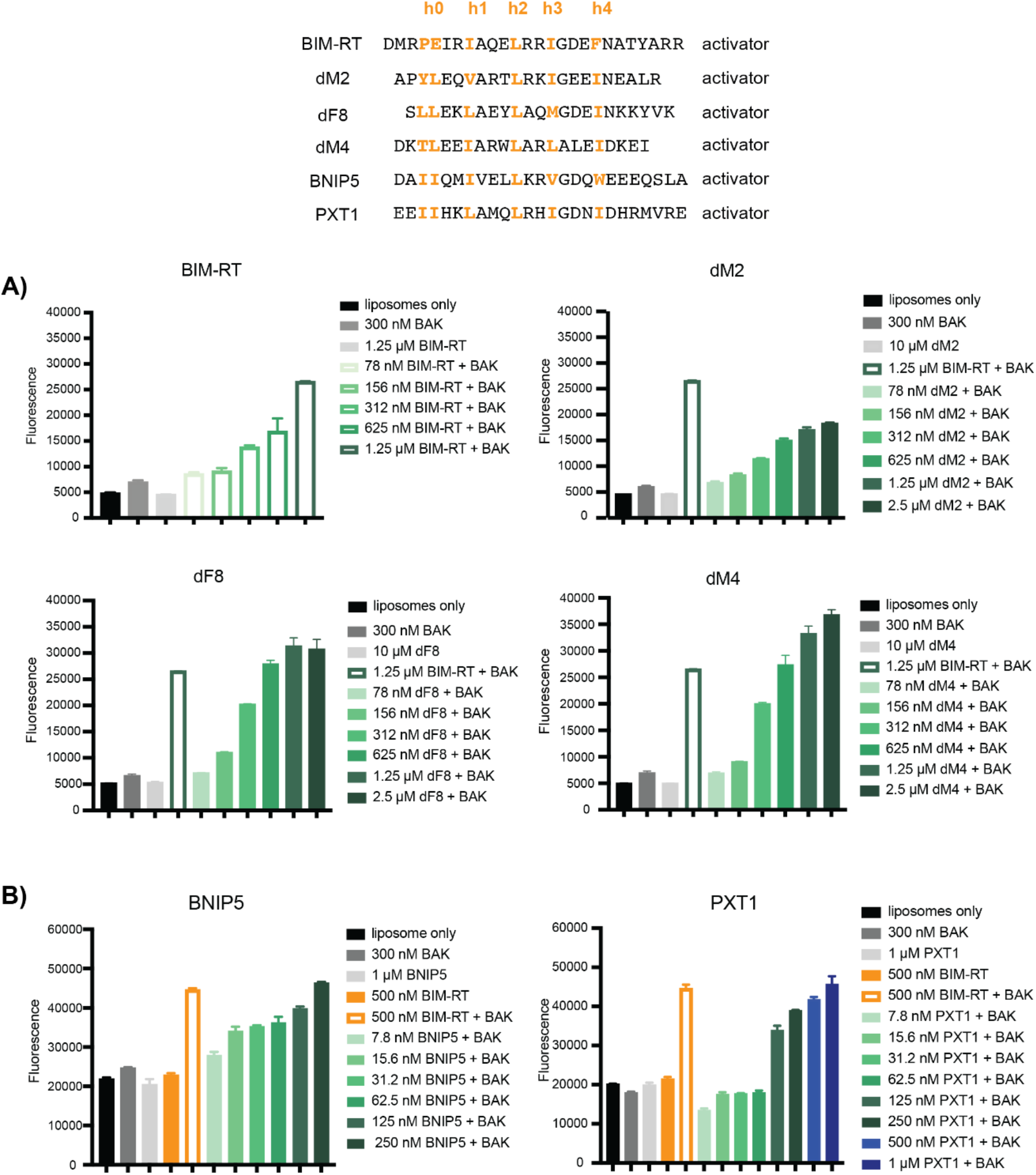
dM2, dF8, dM4, BNIP5, and PXT1 peptides function as BAK activators. Dye-containing liposomes were incubated with 300 nM BAKΔC25-His_6_ (indicated as “BAK” in figure panels) and newly identified BAK-binding peptides. **A)** Synthetic peptide concentrations from 78 nM to 2.5 µM were tested in comparison to 1.25 µM BIM-RT as a positive control (open bars). Plots show fluorescence signal at 1.5 hours. **B)** Peptides from BNIP5 and PXT1 were tested at concentrations ranging from 7.8 nM to 250 nM (BNIP5) or 7.8 nM to 1 µM (PXT1) compared to 500 nM BIM-RT as a positive control (open bars). Plots shows show fluorescence signal at 1 hour. In all panels, error bars indicate standard deviations over three replicates.

Not all BAK-binding peptides functioned as activators in our assay. Specifically, dF2, dF3, dF4, BK3, and dF7 peptides did not result in dye release at 300 nM BAKΔC25-His_6_ with 5 µM (dF3, dF4, and dF2) (**Figure 4A**) or 10 µM (dF7) peptide (**Figure 4B**) (see **Figure S11** for raw data). We postulated that BAK-binding peptides that do not activate may function as inhibitors, so we tested dF2, dF3, dF4, BK3, and dF7 with 300 nM BAKΔC25-His_6_ and varying concentrations of BIM-RT (**Figure 4**). Given the limitations of the *in vitro* assay, including liposome variability from batch to batch as well as a plateau reached at high activation levels (loss of activation resolution), we first picked a concentration of BIM-RT at which approximately half-maximal dye release was observed (200 nM for experiments in **Figure 4A** and 625 nM for experiments in **Figure 4B**). We subsequently conducted experiments with 300 nM BAKΔC25-His_6_ and varying concentrations of candidate inhibitor peptides that were mixed with BIM-RT at the established half-maximal concentration. Peptides dF2, dF3, dF4, BK3, and dF7 showed concentration-dependent inhibition of BAK activation by BIM-RT. dF2 was the most potent inhibitor, with approximately full BAK inhibition reached at 1.25 µM. Computationally designed BK3 gave complete inhibition at 5 µM. We also tested previously published BIMh3PcRT, an inhibitor peptide that contains a non-natural amino acid (Brouwer et al., 2017), and noted that all four of our peptides were more potent inhibitors in this assay.

**Figure 4.**
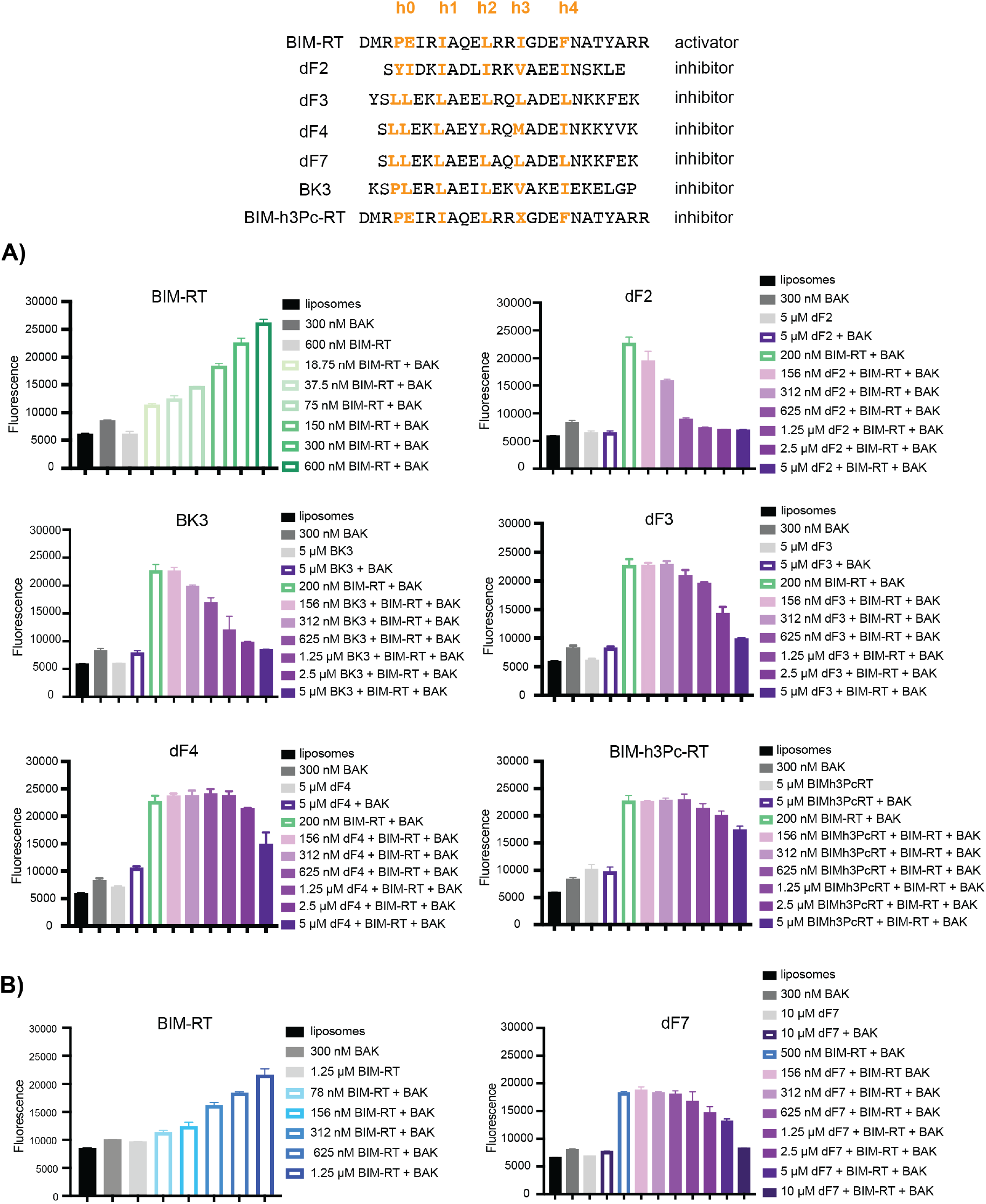
Peptides dF2, dF3, dF4, BK3, BIM-h3Pc-RT, and dF7 inhibit activation of BAK by BIM-RT. Liposomes containing dye were incubated with BIM-RT at a range of concentrations and then with BIM-RT plus candidate peptide inhibitors. **A)** Liposomes with 300 nM BAKΔC25-His_6_ (shown as BAK in the figure) plus 200 nM BIM-RT were incubated with dF2, BK3, dF3, or dF4. Previously published BIM-h3Pc-RT peptide served as a positive control for inhibition (Brouwer et al., 2017). Plots show fluorescence signal at 1.5 hours. **B)** Liposomes with 300 nM BAKΔC25-His_6_ (shown as BAK in figure) plus 500 nM BIM-RT were incubated with dF7 at increasing concentrations. Plots show fluorescence signals at 1.5 hours. In all panels, error bars indicate standard deviations over three replicates. More replicates in progress.

The candidate BH3 peptide from human TRIM58 with measured K_d_ > 20 µM did not function as an activator or inhibitor of BAK. NBEAL2, SLC19A1, and SPNS1 were not soluble in water and were not tested further, whereas CASP3, TRPM7, and TERT showed BAK-independent dye release from liposomes (**Figure S12**).

Amphipathic helical peptides can fold into coiled coils or other helical-assembly structures (Truebestein & Leonard, 2016). To test whether BH3-like peptides that functioned as inhibitors bind to BIM-RT or Y-BID, as well as to BAK, and whether this might contribute to the observed inhibition, we conducted circular dichroism (CD) experiments to test for peptide hetero-association, comparing the helicity of peptides alone vs. in a hetero-mixture (**Table S6 and Figure S13**). The lowest peptide concentration at which we were able to obtain a low-noise CD signal was 15 µM, which is greater than the concentration used in our liposome assay. However, even at this high peptide concentration, no association was observed between any of dF2, dF3, and BK3 and BIM-RT or Y-BID. Previous biophysical and structural evidence has shown that BIM BH3 peptides can self-associate to form tetramers (Assafa et al., 2021). To test whether our peptides contained a significant amount of alpha helical content, possibly due to self-association, we calculated the mean residue ellipticity (MRE) and found that activators and inhibitors both exhibited a range of helicities (**Table S6**). Inhibitor peptides dF2 and dF3 showed 28% and 83% helicity, respectively, while activator peptides BIM-RT and PumaM3dI showed helicities of 5% and 50%, respectively. Given that short peptides typically have very low helicity as monomers, we anticipate that the high-helicity signals at 15 µM likely result from peptide self-assembly, although we did not pursue this further.

### Short peptides from human BNIP5 and PXT1 and non-native designed BH3-like peptides induce BAK-dependent MOMP in cells

Testing for activation function in cells is complicated by the presence of many different BCL-2 family members including BAK and BAX, pro-apoptotic BH3-only proteins, and anti-apoptotic proteins. Inhibiting anti-apoptotic proteins can lead to MOMP through an indirect sensitization mechanism (Chen et al., 2005) (Willis et al., 2007). Nevertheless, to look for evidence of function of our newly discovered activators in cells we used permeabilized, siRNA-treated HeLa cells, which express human BAK and BAX and low levels of anti-apoptotic proteins compared to other cell lines (Placzek et al., 2010). We tested for peptide-induced release of cytochrome c using BH3 profiling, as previously published (Fraser et al., 2019) (**Figure S14, Figure 5, and Table S7**). We compared cytochrome c release across WT, BAK only, BAX only, and BAK/BAX double knockout (DKO) cells using BID, BIM, and PUMA BH3 activating peptides as positive controls and PUMA2A peptide as a negative control (**Table S7 and Figure 5**). Western blots confirmed effective siRNA knockdowns of BAK and BAX (**Figure S15**), and DKO cells were resistant to peptide-induced MOMP for BID, BIM, PUMA. Negative control peptide PUMA2A did not induce MOMP in any cells (**Figure 5**) (Fraser et al., 2019). Using this assay, we compared the mitochondrial functions of new peptides that activate BAK in liposome assays with established activators BIM and BID.

**Figure 5.**
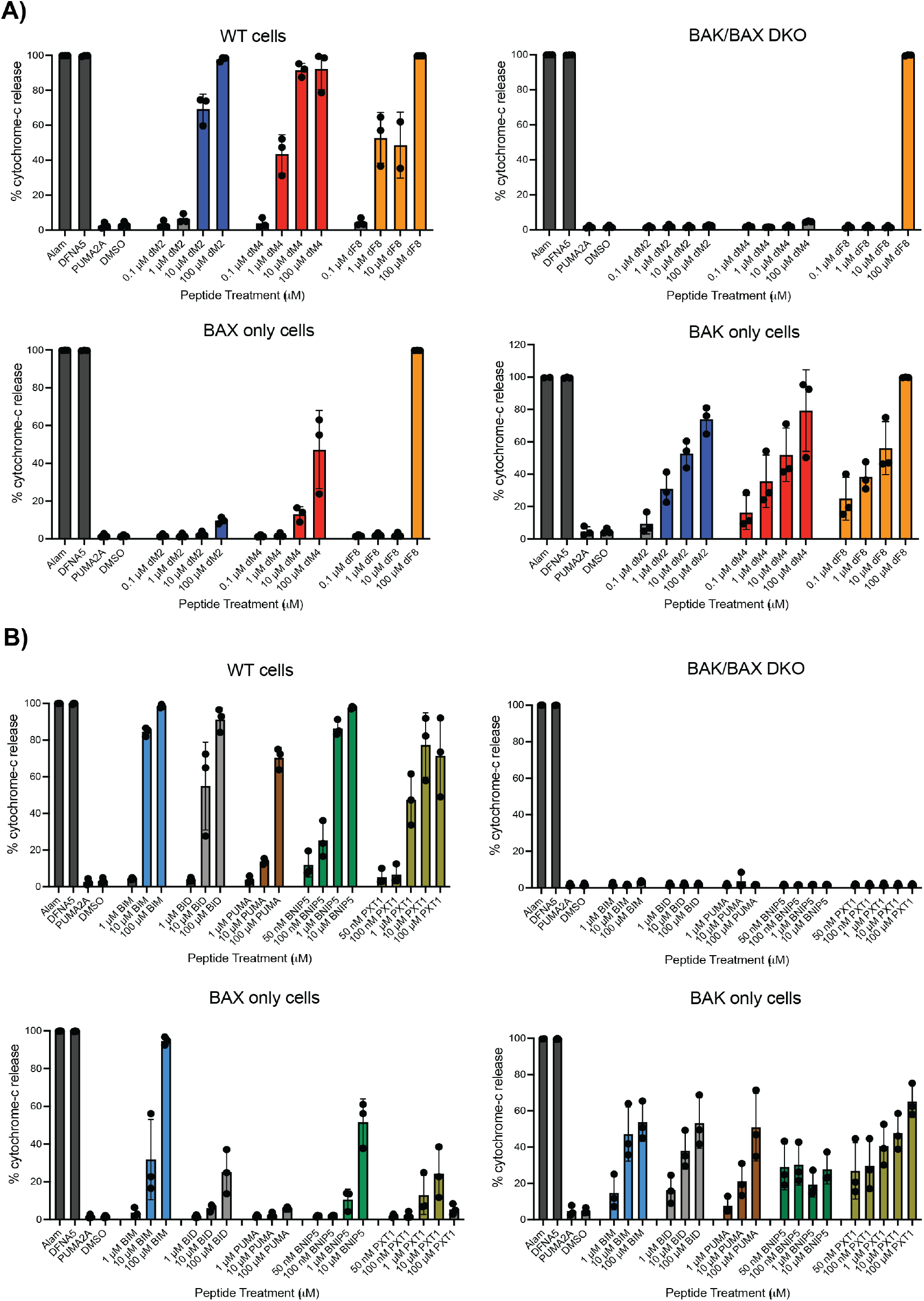
Non-native peptides and human BNIP5 and PXT1 peptides induce membrane permeabilization in cells. Percent cytochrome c release was measured through BH3 profiling in permeabilized HeLa cells including WT, BAK only, BAX only, and BAK/BAX DKO cells. **A)** Non-native peptides dM2 (navy blue), dM4 (red), and dF8 (orange) were tested. **B)** Known activators BIM, BID, and PUMA were tested in addition to BNIP5 and PXT1. Data are from three independent experiments.

We observed that BNIP5, PXT1, dM2, dF8, and dM4 induced the release of cytochrome c, consistent with our results showing that these peptides activate BAK in liposome assays (**Figure 5 and Figure 3**). Peptides dM2, dF8, and dM4 preferentially induced MOMP in the BAX knock-down cells but not the BAK knock-down cells, consistent with selective activation of BAK (**Figure 5A**). Interestingly, BNIP5 induced full cytochrome-c release at 10 µM in WT cells and gave 50% cytochrome-c release at 10 µM in BAX-only cells (**Figure 5B**). Moreover, PXT1 induced 80% and 50% cytochrome-c release in WT and BAK-only cells, respectively.

### Activators and inhibitors engage BAK using similar binding modes

To assess whether differences in binding geometry distinguish activators from inhibitors, we solved crystal structures of examples of each type of peptide bound to monomeric BAK and carried out a systematic comparison of these new structures and other complexes available from prior work (**Figure 6 and Table S8**).

**Figure 6.**
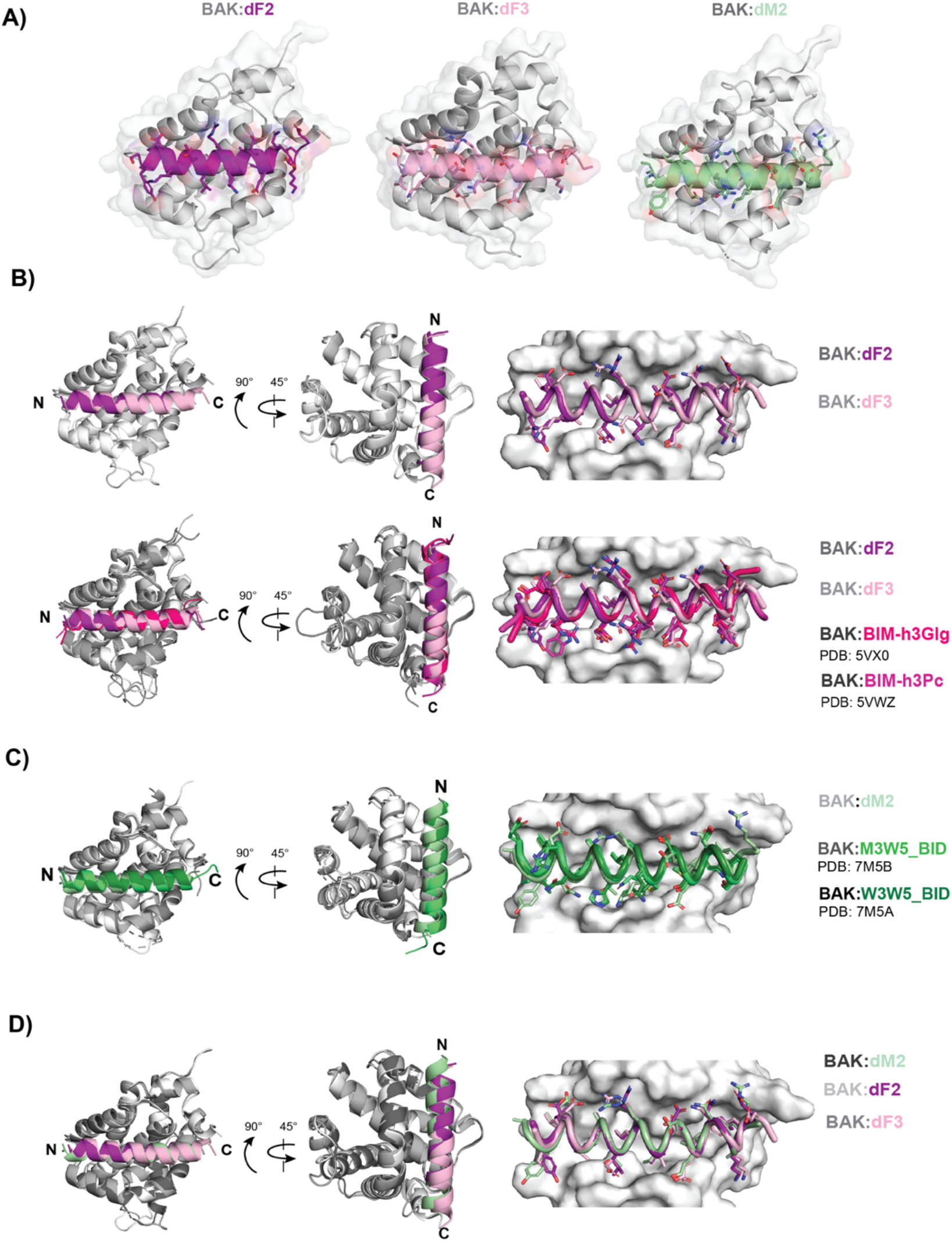
dF2, dF3, and dM2 bind to BAK similarly to other inhibitor and activator BH3 peptides. **A)** Structures of BAK (grey) bound to dM2 (pale green), dF2 (purple), and dF3 (light pink). Inhibitors are shown in shades of purple and pink and activators are shown in shades of green. **B)** (Top) Superimposed representations of BAK bound to dF2 and dF3 inhibitors (left side). BAK-contacting residues in the peptide are shown with sticks (right side). (Bottom) Superposition of BAK bound to inhibitors containing non-natural amino acids including BIM-h3Glg (hot pink, PDB:5VX0) and BIM-h3Pc (light magenta, PDB:5VWZ) compared with newly solved dF2 and dF3 complex structures. **C)** Superposition of BAK complexes including activator peptides: dM2 (light green) and previously published activators M3W5_BID (lime green, PDB:7M5B) and W3W5_BID (forest, PDB:7M5A). **D)** Superimposed cartoon representations of dM2 (activator), dF2 (inhibitor), and dF3 (inhibitor). Further comparisons are included in **Figures S17 and S18**.

To compare inhibitor complexes, we resolved a 1.3 Å structure of BAK (grey) bound to dF2 (purple) and a 1.99 Å structure of BAK bound to dF3 (light pink) (**Figure 6A**). Superposition based on the highly similar conformations of BAK in these two complexes showed that dF2 and dF3 bind very similarly, with no notable differences in the peptide backbone or the positioning of BAK-contacting side chains (**Figure 6B**). BAK complexes with inhibitor peptides BIM-h0-h3Glt, BIM-h3Glg, and BIM-h3Pc-RT, previously solved by Brouwer et al., include both BAK monomers and core-latch domain-swapped BAK dimers bound to peptides that include a non-native amino acid that binds deep in the BAK domain core (Brouwer et al., 2017). The two core-latch dimer complexes, with BAK bound to either BIM-h0-h3Glt (raspberry red) (PDB:5VWX) or BIM-h3Pc-RT (salmon) (PDB:5VWY) differ in sequence at only a single peptide residue and are highly structurally similar, so we chose PDB:5VWY as a representative structure (**Figure S16A**). We compared this core-latch dimer to monomeric BAK bound to BIM-h3Glg (hot pink) (PDB:5VX0) and once again observed no significant differences between the two (**Figure S16B**). Finally, we compared monomeric BAK bound to BIM-h3Glg (hot pink) (PDB: 5VX0) and BIM-h3Pc-RT (light magenta) (PDB:5VWZ) to our novel inhibitors (**Figure 6B**). Overall, we did not see any significant differences in the structure of BAK, nor in the peptide backbone or side-chain arrangements, leading us to conclude that our inhibitors adopt the same binding mode as previously published peptides that contain long, non-natural amino acids that make interactions not accessible to native residues.

To compare our novel activator dM2 to previously published activator complexes, we obtained a 1.3 Å structure of dM2 bound to monomeric BAK (**Figure 6A**). Other structures of activator peptides include mutants of native activator BID such as W3W5_BID (PDB: 7M5A), which contains Trp residues at positions 3d and 4e, and M3W5_BID (PDB:7M5B), with Met and Trp at these positions (**Figure 6C**) (G. Singh et al., 2022). Singh et al. do not classify W3W5_BID as an activating peptide, but we do so here based on data demonstrating that it induces robust activation, similar to that of BID and BIM BH3 peptides, when used at a concentration of 1 µM in a liposome assay (G. Singh et al., 2022). Superposition of BAK bound to dM2 (pale green), W3W5_BID (forest green), and M3W5_BID (lime green) reveals high similarity and excellent structural alignment with no observable distinctions among the three peptides, except two side-chain rotamers (**Figure 6C**).

We also compared BAK-bound activator dM2 with crystal structures of core-latch domain-swapped dimers of BAK bound to BAK BH3 (PDB: 7M5C) and BIM-RT (PDB:5VWV), which are both activating peptides (G. Singh et al., 2022). A comparison of BAK structures indicates that the BAK BH3-bound binding groove opens up less, compared to the BAK dM2 binding groove (**Figure S17A**). A slight divergence at the N-terminus of superimposed dM2 (pale green) and BAK BH3 (deep teal) peptides also leads to differences in polar contacts between the two peptides (**Figure S17A**). Specifically, BAK BH3 forms additional polar contacts with BAK H99 and Q98, while dM2 forms polar contacts with S116 on BAK (**Figure S17A**). Another distinction is observed when superimposing complexes of BAK bound to dM2 (pale green) and BIM-RT (sky blue) (**Figure S18B**). A small difference at the C-terminus of the peptides is associated with a difference in polar contacts. **Figure S18B** shows a top view of dM2 (yellow contacts) and BIM-RT (green contacts) hydrogen bonding networks. The biggest variation is observed at residues E92, Y89, and R88 on BAK that can form interactions with dM2, but not BIM-RT. This gain of contacts, however, does not prevent dM2 from functioning as an activator, although dM2 is a weaker activator compared to BIM-RT at 1.25 µM peptide (**Figure 3**). Overall, we concluded that structures of activators display more differences among them compared to inhibitors, based on structures solved so far.

Finally, to compare structural differences between activators and inhibitors, we compared our three new structures of BAK complexes bound to inhibitors dF2 (purple), dF3 (light pink), and activator dM2 (pale green) (**Figure 6D)**. Interestingly, peptide dM2 engages BAK using the same geometry as the inhibitor peptides dF2 and dF3; it is not distinguished by a unique binding mode nor by any difference in the BAK structure. We extended our analysis by incorporating another inhibitor BAK:BIM-h3Glg (hot pink), and the activators BAK: BAK BH3 (deep teal) and BAK:BIM-RT (sky blue) into this comparison (**Figure S18**). With the exception of BAK:BAK BH3, which is an outlier, the peptides bound very similarly in the groove, with little variation in positioning or axial rotation (**Figure S18A**). Given the previously noted importance of residues at hydrophobic positions, we carefully examined the hydrophobic residues that engage the hydrophobic pockets of BAK and found that, in agreement with the similar binding modes, all activators and inhibitors aligned well at the hydrophobic positions (**Figure S18B**). Singh et al. have reported the presence of an electrostatic network that includes α1 helix and involves residues R42, E46, D90, N86, Y89, R137 in BAK (G. Singh et al., 2022). We found that the BAK:BAK BH3 complex shows differences in this network compared to the 5 other BAK complexes (**Figure S18B**). For example, side chains D90, R137, N86 showed alternative rotamer placements. However, the five related activator and inhibitor structures share a similar network in this region, dissimilar to that observed for BAK:BAK BH3, indicating that this local structure is not a characteristic of activators generally. Overall, our structural analyses did not reveal consistent differences in binding mode or residue interaction networks that could distinguish structures of BAK bound to activators vs. inhibitors.

Other groups have reported the presence of a cavity at the protein-peptide interface of complexes of BAK and BAX bound to BH3 peptides, and it has been suggested that this is important for BH3-induced destabilization and activation (Czabotar et al., 2013) (Brouwer et al., 2017) (G. Singh et al., 2022). The BAK:BIM-RT peptide complex cavity is located between BAK α1, α2, α3, and α5 helices with an estimated volume of 435 Å^3^, while the BAX:BID peptide complex is located between BAX helices α2, α5, and α8 with an estimated volume of 140 Å^3^ (Brouwer et al., 2017) (Czabotar et al., 2013). Inhibitor peptides with non-natural amino acids occupy this cavity in BAK (Brouwer et al., 2017), suggesting that stabilizing this region of BAK may be important for inhibition. We measured differences in cavity size across BAK:peptide complexes for activators and inhibitors and found no clear associations between cavity volume and peptide function (**Figure S19**). The activator BAK:BIM-RT complex (PDB:5VWV) had the largest cavity, with a volume of 404 Å^3^, while the second-largest was found in the inhibitor BAK:dF2 complex, with a cavity volume of 343 Å^3^. Activator complexes BAK:M3W5_BID (PDB: 7M5A) and BAK:BAK BH3 (PDB: 7M5C) showed the smallest cavities, with volumes of 68 Å^3^ and 104 Å^3^, respectively.

We generated peptide sequence logos to compare activators and inhibitors (**Figures S20**). Sequence logos showed that both inhibitors and activators display the canonical L-x(3)-G/A-D/E motif in addition to a preference for hydrophobic residues at the BAK hydrophobic pocket positions. However, activators showed more sequence variability at the site that binds the h0 pocket. For example, a negatively charged glutamate is present at position 2a in activator BIM, but not in any of the other activators. Furthermore, inhibitors appear to show a strong preference at positions 2b, 2c, 2f for negatively, positively, and negatively charged residues, respectively. However, an inspection of BAK:peptide complex structures shows that these residues are solvent exposed and do not form significant contacts with BAK. Similarly, lysine residues in inhibitors at positions 4c do not make polar contacts with BAK. Lastly, both activator and inhibitor peptides contain a negatively charged glutamate at position 3g that forms a salt bridge with arginine 88 on BAK. Overall, we did not observe any consistent differences in peptide sequences that were indicative of activator or inhibitor function.

Finally, we computed pairwise RMSD values between peptides in BAK:peptide complexes including inhibitors dF2, dF3, and BIM-h3Glg and activators dM2 and BIM-RT (**Figure S21**). We did not include the BAK:BAK BH3 complex (PDB:7M5C) in our analysis, given that this structure contains clear differences in the BAK groove and is a structural outlier in this set of complexes. Our pairwise comparisons of activator vs. inhibitor peptides gave an average RMSD value of 1.2 Å and did not indicate any significant distinction between the two groups.

### Binding affinities and kinetics do not distinguish activators and inhibitors of BAK

To test whether activator vs. inhibitor peptides bind to BAK with systematically different affinities or kinetics, we conducted experiments using bio-layer interferometry (BLI) (**Figure 7**). Briefly, the biotinylated protein b-His_6_-BAK_16-186_ was purified and immobilized on streptavidin coated tips, and binding and dissociation rates of His_6_-SUMO-peptide fusions were measured. We examined trends in affinities, k_on_, and k_off_ values for 5 inhibitors and 3 activators (**Figure 7** and **Table S9**). We were unable to measure BLI affinities for activators dM4 and PXT1, given their weak binding and fast kinetics. We also measured peptide affinities using fluorescence anisotropy (**Table S10**), and the observed trends were in agreement with the BLI measurements (**Table S11**). Although in general the activators were weaker binders compared to inhibitors, this was not consistent across all peptides. For example, inhibitors dF4, dF3, dF2, and BK3 bound more tightly compared to activators dF8 and dM2, but the high affinity activator BNIP5 BH3 was an exception to this trend.

**Figure 7.**
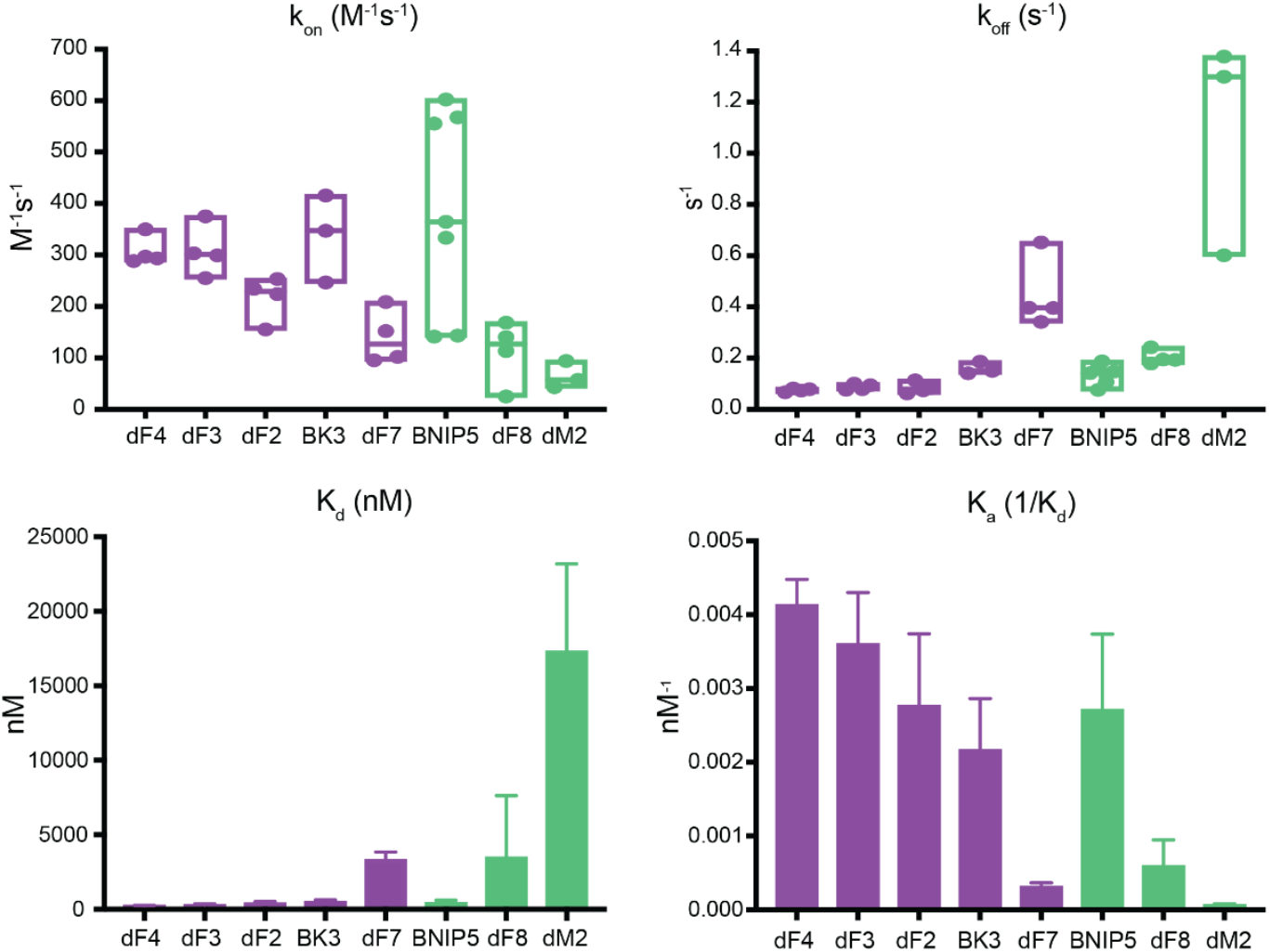
Neither rate constants nor affinities are indicative of peptide activator vs. inhibitor function. Replicates consisted of dF4 n=4, dF3 n=4, dF2 n=4, BK3 n=3, dF7 n=4, BNIP5 n=7, dF8 n=4, and dM2 n=3. Inhibitors are in purple and activators are in green. Biotinylated b-His_6_-BAK_16-186_ was immobilized on streptavidin-coated tips and tested for binding to recombinantly expressed and purified His_6_-SUMO-peptides. At least three replicate measurements for each peptide are indicated as points on the box plots. K_d_ values were computed as the ratio of k_off_/k_on_.

Our results did not reveal consistent differences in the binding kinetics of activators and inhibitors. For example, we found that activators dF8 and BNIP5 had dissociation rate constants similar to that of inhibitor BK3. Activators dF8 and dM2 showed slower association compared to inhibitors dF4, dF3, dF2, and BK3; but inhibitor dF7 bound with similar kinetics as activator dF8. Activator dM2 displayed fast binding kinetics, making it difficult to determine k_on_ with confidence, even at low peptide concentrations. In conclusion, 4 out of 5 inhibitors showed higher affinities compared to 2 out of 3 activators, although the number of activators for which we could get reproducible data was smaller than that of inhibitors (Singh et al., 2022). Measurements for other peptides, reported in the literature, further support the absence of an affinity requirement for activation or inhibition, as discussed below.

## DISCUSSION

Pro-apoptotic BAK is a key regulator that directly disrupts the mitochondrial outer membrane upon cell death stimulus (Chittenden et al., 1995). This irreversible step is regulated by BH3-only proteins binding to membrane localized BAK, which triggers a series of conformational changes that lead to BAK homodimerization and subsequent higher order cluster formation and MOMP (Kim et al., 2009). We sought to elucidate features that allow some, but not all, BH3-only proteins to activate pro-apoptotic BAK. To further decipher the complex activation mechanism that governs BAK function, we focused our studies on the initial binding event that triggers the cascade of subsequent conformational changes.

Seeking a broad panel of BAK binders, we used peptide screening and computational design to discover 19 non-native peptide binders of BAK. These peptides displayed a range of profiles in yeast surface-display experiments, and we tested eight peptides with strong binding signals in a liposome assay that serves as a proxy for MOMP. We found that five peptides functioned as inhibitors and three functioned as activators of BAK and led to liposome permeabilization. To our knowledge, previous groups have only tested point or pair mutations of previously known activators. Our results dramatically expand the sequence space that has been tested for BAK-regulating function and reveals that this space includes both activators and inhibitors that could serve as the basis for developing BAK-modulating therapeutics. Notably, inhibitor peptides BK3, DF2, and DF3 are more potent in liposome assays than the previously reported BAK-inhibitor peptide BIM-h3Pc-RT yet are composed entirely of native amino acids (Brouwer et al., 2017). However, we did not compare our peptides to other tighter variations of BIM-h3Pc-RT containing other non-natural amino acids (NNAs) at the 3d position engaging with the BAK h3 pocket.

To our knowledge, only five native BH3 peptides are consistently reported to bind and activate BAK including BID, BIM, PUMA, the BH3 sequence within BAK itself, and the corresponding region in BAX (Moldoveanu et al., 2013) (Sarosiek et al., 2013) (Kim et al., 2009) (Hockings et al., 2015) (Llambi et al., 2011). In this work, we discovered nine previously unidentified human proteins that contain BH3-like regions that directly interact with BAK, most of which are predicted to be structurally accessible in the context of the full-length protein **(Table S4)**, and two of which showed potent induction of MOMP in cells. BNIP5 (also referred to as C6orf222) and PXT1 bound BAK with different affinities (nanomolar vs. low micromolar) but both induced BAK-dependent membrane permeabilization in cells. BNIP5 and PXT1 were more potent than BID and PUMA in this profiling assay. Also, BNIP5 was more potent than PXT1 in WT cells. Interestingly, BNIP5 and PXT1 retained potency at 10 µM in BAK depleted cells, indicating that both induce BAX-mediated MOMP. Both BNIP5 and PXT1 bind all five anti-apoptotic BCL-2-family proteins with nanomolar affinities (DeBartolo et al., 2014), which means that the function of these proteins as activators vs. sensitizers likely depends on the relative expression levels and localization of pro-vs. anti-apoptotic proteins. BNIP5 has unknown function, though transcript levels are high in the colon, small intestine, pancreas, and stomach as characterized in hematopoietic cells (Luck et al., 2020). Peroxisomal testis-specific 1 (PXT1) is expressed in male germ cells, where overexpression induces apoptosis of spermatocytes (Kaczmarek et al., 2011). Our work indicates that BNIP5 and PXT1 are two additional BH3-only activator proteins that may have unexplored biological implications in the apoptotic regulatory network.

The features that make a peptide an activator vs. an inhibitor of BAK are not known. Prior to our study, others had shown that a single-residue substitution at a BAK-binding hydrophobic position could convert an activator to an inhibitor (Brouwer et al., 2017), suggesting that minimal sequence differences are sufficient to alter BAK function. Furthermore, consistent with a previous report for BAX (Czabotar et al., 2013), we found differences in activation potency among BH3-only peptides that establish that an amphipathic peptide with a BH3 motif including L-x(3)-G/A-D/E (x = any amino acid) is not sufficient to activate BAK. Single-residue substitutions in BH3 peptides can increase BAK activation activity and convert a non-activator (NOXA) to a weak activator and a moderate activator to a more potent activator (PUMA), as has previously been observed for BAX. As-yet unknown sequence and structural features drive varying functional outcomes.

A possible mechanistic explanation for activation could be that activators vs. inhibitors engage BAK with different binding modes, in distinct geometries. Dissociation of the BAK “latch” (α6-α8) from the core (α2-α5) is thought to be an early step in activation, and it is plausible that some peptides bind in a way that induces a conformational change that is propagated to the latch via a pathway of allosteric communication. Supporting the possibility of such a conformational change, Singh et al. showed a BAK BH3 peptide docks into the binding groove of a BAK monomer in a distinct pose that is accompanied by a rearrangement of a BAK salt-bridging network (G. Singh et al., 2022). However, our analysis of multiple crystal structures of BAK-peptide complexes, including three structures that we solved in this work, showed that 3 inhibitors and 3 activators bind with very similar geometry and contacts; peptides in these two functional groups cannot be structurally distinguished by any criterion that we could discern. The re-arranged BAK residues in the BAK:BAK BH3 complex appear to be specific to that interaction, and not characteristic of activators more broadly.

Considering other mechanisms, we reasoned activators vs. inhibitors might differ in their binding kinetics. The association of activators with BAK is transient, and the groove in which activator and inhibitor BH3 peptides bind re-shapes to form the groove that is occupied by the BH3 helix of a partner BAK molecule in the so-called “BH3:groove homodimer” that is critical for membrane poration. Following BH3 binding that induces a change in BAK, which may correspond to core-latch dissociation, dimerization requires activator peptide dissociation followed by a rearrangement in which two neighboring BAK monomers exchange BH3 helices. Peptide dissociation may therefore set up a competition between activator or inhibitor BH3 peptide re-binding vs. BAK dimerization via BH3 exchange. In this model, peptides that re-bind more slowly than BAK dimerization would function as activators, whereas those that re-bind quickly and continue to occupy the canonical groove would function as inhibitors. Although we found that most of the activator peptides in our study bound more slowly to BAK than did most of the inhibitors, this was not consistent across all peptides. Specifically, activator BNIP5 had the greatest k_on_ rate constant (5.8 × 10^−4^ M^-1^s^-1^) compared to the 5 inhibitors and 3 activators that were tested. Furthermore, inhibitors did not necessarily dissociate more slowly. For example, inhibitor dF7 has a greater k_off_ rate constant (0.4 s^-1^) compared to activators BNIP5 (0.16 s^-1^) and dF8 (0.2 s^-1^). Overall, our kinetic data indicate that k_on_ and k_off_ rate constants measured using bio-layer interferometry do not determine whether or not a peptide activates or inhibits BAK.

Another possibility is that activators and inhibitors vary in their affinity for BAK. This model is supported by the increase in BAK-binding affinity of BIM when position 3d is substituted with a pentyl-carboxylate (h3Pc) and its variants Glg and Glt. These substitutions increase the affinity of BIM to K_d_ = 1.3 µM, 21 µM or 1 µM, relative to weakly binding BIM (with an affinity that is too weak to measure using BLI), and also convert BIM from an activator to an inhibitor (Brouwer et al., 2017). Peptides that bind tightly to BAK monomers and stabilize that inactive state would be expected to act as inhibitors. This is the general trend that is observed in our data (**Figure 7, Table S11**), but there are exceptions. For example, BNIP5 is an activator and tight binder, with an affinity of 424 nM. A more recent study also shows that tight-binding peptides M(3)W(5) BID BH3 (K_d_ = 690 nM) and M(3)W(5) BID BH3 (K_d_ = 250 nM) can activate BAK in liposomes at 1 µM (G. Singh et al., 2022). Leshchiner et al. have also shown that stapled BID SAHB peptides with low nanomolar affinities for binding to murine full-length BAK activate in liposome-based assays (Leshchiner et al., 2013).

We propose a modification of the affinity model that reconciles all of our observations as well as other data in the literature. In **Figure 8**, we use a free energy landscape diagram to illustrate how BH3 peptides may act as *catalysts* of activation by binding differentially to the BAK ground-state monomer and the transition state for BAK activation. In the figure, we represent BAK tethered to the mitochondrial outer membrane as an inactive monomer. Following activation, BAK forms the lower energy BH3:groove homodimer (PDB: 7K02). Between the monomer and the dimer, on this pathway, BAK must pass through a transition state of unknown structure. In the context of this free energy landscape model, activators and inhibitors differ in their affinities for distinct BAK conformational states. As for enzyme-catalyzed reactions, tighter binding of a BH3 peptide to the activation transition state vs. the monomer ground state will lower the energy barrier and promote activation. In contrast, tighter binding to the ground state than to the transition state will inhibit activation.

**Figure 8.**
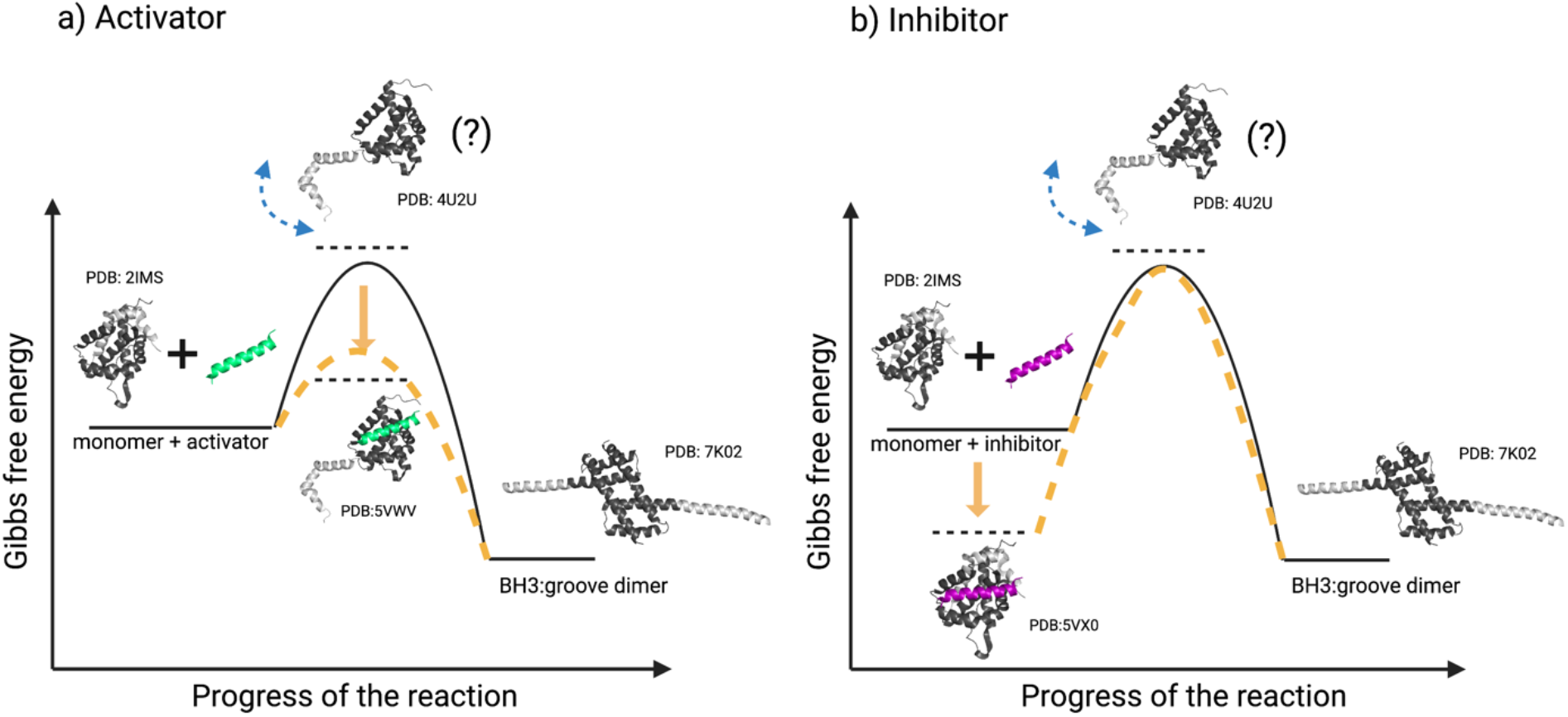
Free energy diagram for BH3 peptide activation or inhibition of BAK. Crystal structures of the BAK monomer and putative intermediates are used to illustrate steps in activation, with the BAK core in dark grey and the latch in light grey. Monomeric BAK (PDB: 2IMS) is placed at a higher energy compared to the BH3:groove homodimer (PDB:7K02), which is presumed to be embedded in the outer mitochondrial membrane (not shown). The structure or nature of the transition state is not known, and is labeled with a question mark, but may resemble an unlatched conformation of BAK as discussed in the text (here represented using PDB:4U2U). **A)** Illustrates stabilization of a putative transition state of BAK by binding of an activator (green). **B)** Illustrates stabilization of monomeric BAK in the presence of an inhibitor (purple).

Multiple conformational changes of BAK need to occur to reach the dimeric state, including core-latch dissociation and dimerization via exchange of BH3 helices. Several groups have reported exposure of the BH3 region as an indicator of activation (Moldoveanu et al., 2006). Because existing structures of domain-swapped dimers with dissociated latch regions do not exhibit any rearrangements of the BH3 region, we assume that this step follows latch dissociation, as illustrated in **Figure 8**. Existing data do not establish which conformational change corresponds to the rate-limiting step. However, all of our peptides – both activators and inhibitors – can bind to the monomer state, and our model requires that at least the activators also bind to the transition state. Activators and inhibitors are similar in sequence and structure, and peptides that differ by just two mutations can differ in their function (inhibitor dF4 and activator dF8). These arguments suggest that the monomer and the transition state share structural similarities and therefore that the transition state is “early” in the pathway leading to dimer formation. For this reason, and because we expect there to be a large energy cost for latch dissociation that disrupts stabilizing intradomain interactions, we speculate that disengagement of the latch from the core is the rate-limiting step, and we use the core-latch dissociated structure (PDB: 4U2U) as a model for the transition state in **Figure 8** and **Figure S22**. However, our data do not rule out other scenarios – such as rate-limiting dissociation of the BH3 helix from the BAK core – which can also be considered.

Notably, it is easy to incorporate the effects of many peptides, and even non-peptide binders into this model. BIM peptide variants with non-natural amino acids at position 3d stabilize the monomeric ground state more than the transition state, whereas stapled BID SAHB peptides with low nanomolar affinities for murine full-length BAK stabilize the transition state in preference to the monomer (Brouwer et al., 2017) (Leshchiner et al., 2013). In addition to binding the canonical hydrophobic groove, it is possible to bind other regions within BAK and induce activation. For example, antibody 7D10 triggers BAK activation by binding the α1-α2 loop. In the context of our model, this implies that the structure of this region differs between the monomer state and the transition state, and that antibody binding preferentially stabilizes the transition state, lowering the energy barrier to the conformational changes that are required for MOMP to occur.

Our finding that just a few mutations in BH3-like helices can give rise to BAK activators would seem to pose a risk that evolutionary drift in BH3 sequences might lead to unregulated cell death. However, anti-apoptotic BCL-2 members play a key role in restraining activation, by sequestering the exposed BH3 region of pro-apoptotic members BAK and BAX at some point after core unlatching. Nanomolar affinities of the BAK BH3 region for anti-apoptotic BCL-2 family members (K_d_ = 53 nM for BCL-x_L_ and 20 nM for MCL-1) allow for an affinity buffer that can tolerate such mutations. That is, mutated BH3-like helices must bind both BAK and the partner anti-apoptotic protein with an affinity greater than that of the BAK BH3 region in order to induce MOMP. Furthermore, the role of mitochondrial membrane channel VDAC2 in restraining BAK must also be taken into consideration (Yuan et al., 2021).

Overall, we discovered two new human and eight non-native peptide binders of BAK with diverse sequences and functions. These include human proteins BNIP5 and PXT1, which activate BAK in cells. The result suggests the possibility of other BH3-like activators in the human proteome that might contribute to apoptotic regulation. We solved three crystal structures of BAK:peptide complexes including two inhibitors and one activator found that, surprisingly, the binding mode of both inhibitors is highly similar to that of the activator, despite their different functions. In addition, our kinetic data shows that neither peptide binding kinetics nor affinity for BAK is sufficient to distinguish activators from inhibitors. This work highlights the complexities associated with binding to the BAK BH3-binding groove and the resulting changes that occur to regulate BAK function. We speculate on the energetic requirements for peptide activation vs. inhibition functions and propose a model to summarize our findings.

## METHODS

### Peptide synthesis, purification, and concentration determination

Peptides were synthesized in the Swanson Biotechnology Center Biopolymers and Proteomics Core. All peptides were amidated at the C terminus; peptides used for fluorescence assays were labeled at the N terminus using a 5-Carboxyfluorescein, single isomer (5-FAM); peptides used for liposomes and crystallography were acetylated at the N terminus. Non-natural amino acid Fmoc-AsuOtbu-OH was purchased from ChemImpex. Crude peptides were purified by HPLC on a C18 column using a linear gradient of water/acetonitrile and masses were verified using MALDI mass spectrometry. Synthesized peptides used in liposome assays were dissolved in water. Absorbance at 280 nm (A_280_) was measured in Edelhoch buffer (7 M guanidine-HCl + 0.1 M potassium phosphate buffer pH 7.4) using a NanoDrop UV spectrophotometer and then peptide concentration (C) was determined based on Beer’s law with the extinction coefficient calculated based on the number of tyrosines (1490 M^-1^cm^-1^) and tryptophans (5500 M^-1^s^-1^) in the peptide using the ProtParam tool in ExPasy https://web.expasy.org/protparam/. Synthesized peptides used in cell-based assays were dissolved in DMSO. Peptide concentration was also determined using Beer’s Law, however, extinction coefficients were re-calculated based on controls tested in DMSO, giving values of 2287 M^-1^s^-1^ and 6650 M^-1^s^-1^ for tyrosine and tryptophan, respectively.

### Fluorescence anisotropy binding assays

Fluorescence anisotropy assays were performed in 25 mM Tris, 50 mM NaCl, 1 mM EDTA, 20 mM HEPES at pH 8.0, with 5% DMSO (FP buffer), in Corning 96-well, black, polystyrene, nonbinding surface plates. Two-fold serial dilution of b-His_6_-BAK_16-186_ to generate 24 points was done in Eppendorf tubes and then protein solutions were transferred to plates. A solution of 100 nM peptide in 50% DMSO and 50% FP buffer was prepared, and 5 µl of this mixture was added to each well to give a final concentration of 10 nM peptide in 5% DMSO. Plates were incubated for 2 hours at room temperature to reach equilibrium and subsequently read at 25 °C in a SpectraMax M5 Multi-Mode microplate reader. Five replicate titrations were performed for each peptide, and the dissociation constant was estimated by fitting each curve to a 1:1 binding model as in Roehrl *et al*. using Python (Roehrl et al., 2004).

### Yeast surface-display

Peptides in **Table S1** were constructed as described in Frappier et al. (Frappier et al., 2019). Briefly, synthetic DNA encoding peptides was amplified and cloned into the yeast surface display plasmid pCTCON2 between Xho1 and Nhe1 restriction digest sites using homologous recombination (Chao et al., 2006). The construct contained a carboxy-terminal FLAG tag after the peptide. Sequences were transformed into yeast strain EBY100 using a Frozen-EZ yeast transformation II kit. Sequence-verified yeast clones were stored in glucose media SD + CAA media (5 g/L casamino acids, 1.7 g/L yeast nitrogen base, 5.3 g/L ammonium sulfate, 10.2 g/L Na_2_HPO_4_-•7H_2_O and 8.6 g/L NaH_2_PO_4_-•H_2_O, 2% glucose) + 20% glycerol. To induce the expression of peptides on yeast, the previously published protocol of Reich *et al*. was adapted (Reich et al., 2016). Plated cells were grown overnight in 5 ml of SD + CAA media at 30 °C. Cells were then diluted to OD600 of 0.05 in 5 ml of SD + CAA media and grown for approximately 8 hours at 30 °C. Once again, cells were diluted to an OD600 of 0.005 and left to grow until reaching an OD of 0.1-0.4. To induce peptide expression, cells were diluted to an OD600 of 0.025 in 5 ml of galactose media (SG + CAA media: 5 g/L casamino acids, 1.7 g/L yeast nitrogen base, 5.3 g/L ammonium sulfate, 10.2 g/L Na2HPO4-•7H2O and 8.6 g/L NaH2PO4-•H2O, 2% galactose) and grown to an OD600 of 0.2–0.5 (approx. 20 hours) at 30 °C.

Peptide-displaying yeast cells were prepared for sorting in low protein-binding 96-well 0.45 μm filter plates. A volume of 50 µl per well at a concentration of 1*107 cells/ml was used. Based on an estimate that an OD600 = 1 corresponds to 3*107 cells/ml, the required volume of cells was pelleted at 14,000 g for 5 minutes. Supernatant was aspirated and cells were washed twice in BSS buffer (50 mM Tris, 100 mM NaCl, 1 mg/ml BSA, pH 8.0). Cells were resuspended to a final density of 1*10^7^ cells/ml in BSS buffer. Then 50 µl of cells per well plus a final concentration of 2.4 µM of pre-tetramerized BAK monomer (see protein expression and purification section) were incubated for 1.5 -2 hours at room temperature (∼25 °C). Cells were filtered with a vacuum and washed twice with pre-chilled BSS buffer. 20 µl of primary anti-HA antibody (mouse, Roche, Indianapolis, IN, RRID:AB_514505) diluted 1:100 was added to the mixture and incubated for 15 minutes at 4 °C in BSS buffer. Cells were filtered and washed twice with BSS buffer. Next, 20 µl of APC-conjugated secondary antibody (rat anti-mouse; CD45 Clone 30-F11, RUO from BrandBD Pharmingen) was added at 1:40 dilution and incubated at 4 °C for 15 minutes. Cells were filtered and washed twice with BSS. Finally, cells were resuspended in BSS and transferred to a second 96-well plate for fluorescence activated cell sorting (FACS) and analyzed for BAK binding using a FACSCanto II HTS-1.

### Liposome assay

Large unilamellar vesicles (LUVs) mimicking the outer mitochondrial membrane were made using Avanti Polar Lipids, Inc. lipids dissolved in chloroform including 1-palmitoyl-2-oleoyl-glycero-3-phosphocholine (Avanti: 850457C 16:0-18:1 PC (POPC)), 1,2-dioleoyl-sn-glycero-3-phosphoethanolamine (Avanti: 850725C 18:1 (Δ9-Cis) PE (DOPE)), L-α-phosphatidylinositol (Liver, Bovine) (sodium salt) (Avanti: 840042C Liver PI), 1,2-dioleoyl-sn-glycero-3-phospho-L-serine (sodium salt) (Avanti: 840035C 18:1 PS (DOPS)), 1,2-dioleoyl-sn-glycero-3-[(N-(5-amino-1-carboxypentyl)iminodiacetic acid)succinyl] (nickel salt) (Avanti: 790404C 18:1 DGS-NTA(Ni)), 1’,3’-bis[1,2-dioleoyl-sn-glycero-3-phospho]-glycerol (sodium salt) (Avanti: 710335C 18:1 Cardiolipin) at a (43:27:11:10:4:5) molar ratio. The lipid mixture was dispensed into borosilicate glass test tubes (13 × 100 mm, Fisherbrand) in 15 mg aliquots. Lipid films were made by evaporating the chloroform in the tubes with nitrogen gas using a glass Pasteur pipet in a fume hood. The film was further dried overnight in a flask connected to a vacuum pump. Lipid films in tubes were individually sealed in plastic pouches filled with nitrogen gas and kept in the freezer at -80 °C.

To form liposomes encapsulating dye molecules, 14 mg of 8-aminonaphthalene-1,3,6-trisulfonic acid, disodium salt (ANTS) dye (Biotium #90010) and 40 mg of p-xylene-bis(N-pyridinium bromide) DPX fluorescent quencher (Biotium #80012) were added to a test tube containing lipid film and hydrated with 500 µl of buffer A (200 mM KCl, 10 mM HEPES, 1 mM MgCl_2_ pH 7.5). The tube was vortexed for 1 minute to completely dissolve the lipid film. The solution was then transferred to an Eppendorf tube. The lipid/dye emulsion was then subjected to 17 freeze-thaw cycles using liquid nitrogen and room temperature water. Subsequently, the resulting multilamellar vesicles were extruded up to 21 times through two sheets of polycarbonate membrane and a filter with 100-nm sized pores (Avanti Polar Lipids), until the solution could be pushed from one extrusion syringe to the other without much resistance. The resulting liposomes were separated from free ANTS/DPX dye using a disposable Sephadex G-25 in PD-10 desalting column (GE Healthcare 17-0851-01) using buffer A to flow the solution through the column. The resulting solution was collected manually in Eppendorf tubes every 10 drops. Translucent fractions were selected for dynamic light scattering (DLS) to verify the size of the liposomes. Fractions corresponding to average distributions of 100 nm diameter particles were subsequently used for experiments. Liposomes were kept covered and at 4 °C until use. Liposomes were made fresh for all experiments performed that same day.

The liposome activation assay was performed in a Corning 384-well plate (#CLS3573). Briefly, 30 µl of buffer A, 10 µl of liposomes, 5 µl of peptide, 5 µl of BAK ΔN22 ΔC25 C166S with a C-terminal His_6_ tag (BAKΔC25-His_6_) were added, in that order. The final volume in each well was 50 µl. Peptide and BAKΔC25-His_6_ were added at 10x the desired final concentration in the well. 0.2% Triton X-100, added without peptide or BAK protein, was used as a positive control for dye release. The plate was immediately put in either a Tecan Spark Multimode or a SpectraMax M5 Multi-Mode microplate reader and read at room temperature for 1-2 hours.

### Protein expression and purification

b-His_6_-BAK_16-186_ was cloned into vector pDW363 and transformed into BL21 (DE3) E. coli cells. 8 L of cells were grown in TB media with ampicillin. Absorbance was monitored until an OD600 of 0.6 was reached, at which point 15 mg of biotin and 1 M IPTG (final concentration 1 mM) were added. Cells were then transferred to 16 °C for overnight growth. Cells were spun down the next morning at 7000 g for 15 minutes. Pellets were transferred to falcon tubes and kept on ice. 25 mL of Ni-NTA binding buffer (20 mM Tris, 500 mM NaCl, 5 mM imidazole, pH 8.0) per liter of growth and 50 µL of PMSF at 100 mM to a final concentration of 0.2 mM per L of growth were added. Pellets were resuspended by pipetting up and down and vortexing and subsequently sonicated for a total of 3 minutes of 20 s on/20 s off at ∼60 power duty cycle ∼5 output control. Lysed cell cultures were centrifuged at 12,000 rpm for 30 mins and filtered through a 0.22 µm filter. Ni-NTA resin washed with Ni-NTA binding buffer was incubated with filtered supernatant for 1 hour at 4 °C. Protein-bound resin was poured through a disposable column and washed three times with Ni-NTA binding buffer. BAK protein was eluted with Ni-NTA elution buffer (20 mM Tris, 500 mM NaCl, and 300 mM imidazole, pH 8.0). Eluted sample was purified by gel filtration using a Superdex 75 26/60 column in TBS buffer pH 7.5. For yeast surface display, b-His_6_-BAK_16-186_ was pre-tetramerized by incubating with streptavidin conjugated to R-phycoerythrin (SAPE) (Thermo Fisher Scientific #S866 at 1 mg/ml concentration) at a 4 BAK:1 SAPE molar ratio and incubated on ice for 15 minutes, shielded from light.

His_6_-SUMO-peptide: Peptides fused to SUMO domains contained a His_6_ tag and short flexible linker sequences: His_6_-GSGSG-yeastSUMO-GSGSGSG-peptide sequence. Peptides were expressed and purified as described for b-His_6_-BAK_16-186_.

BAK ΔN22 ΔC25 C166S and BAKΔC25-His_6_ constructs in pGEX 6P3 and pTYB1 vectors, respectively, that encoded expression as a fusion to glutathione S-transferase (GST) were obtained from Peter Czabotar, Walter and Eliza Hall Institute of Medical Research. The plasmid was transformed into BL21 (DE3) *E. coli* cells and grown in 4 L of TB media plus ampicillin and induced with IPTG (final concentration 1 mM) upon reaching an OD600 of 0.5-1.0. Cells were grown overnight at 16 °C. The next morning, cells were lysed and centrifuged at 7000 g for 15 mins. Pellets were transferred to falcon tubes and kept on ice. 25 mL of cold GST buffer (100 mM Tris, 50 mM NaCl, 1 mM EDTA, pH 7.5) per liter of growth and 50 µL of 100 mM PMSF were added per liter of growth media. Pellets were resuspended by pipetting up and down and vortexing and subsequently sonicated for 3 mins total -20s on/20s off at ∼60 power duty cycle ∼5 output control. Lysed cell cultures were centrifuged at 12,000 rpm for 30 min and filtered through a 0.22 µm filter. A bed volume of 5 ml of glutathione resin (GenScript L00207) was used, washed with 50 ml of cold PBS buffer at 4 °C. Supernatant was added to the column, maintaining a flow rate of 10-15 cm/hr. The column was washed with 20x bed volume of GST buffer. The GST tag was cleaved with 200 units of PreScission protease (GE Healthcare, 27-0843-0) for 2 days on the column. Eluted cleaved protein was collected and further purified by gel filtration, using a Superdex 75 26/60 column in TBS buffer pH 7.5.

### Bio-layer Interferometry

Bio-layer interferometry (BLI) experiments were performed on a Sartorius Octet RED96 instrument using streptavidin tips. b-His_6_-BAK_16-186_ in freshly made BLI buffer (PBS buffer (137 mM NaCl, 2.7 mM KCl, 10 mM Na_2_HPO_4_, 1.8 mM KH_2_PO_4_,) plus 0.005% Tween-20, 1% BSA, 2% DMSO, pH 7.5) was immobilized on streptavidin tips until a signal of around 0.6 nm was reached. Control His_6_-tagged SUMO-peptide constructs were subtracted from the raw data using the BLI analysis software.

For kinetic analysis, curves for the dissociation portion of the curve only were fit to a One Phase Decay model using PRISM:

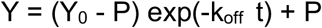

Where t is time in seconds, Y is the receptor binding signal, Y_0_ is Y at time 0 (the initial maximal binding signal that decreases to the plateau signal P), and k is the dissociation rate constant in units of s^-1^. The dissociation constant k_off_ was constrained to be a single value for all peptide concentrations. Given the fit value for k_off_, k_on_ was obtained by fitting the association portion of the data to an exponential equation using a global fit in PRISM:

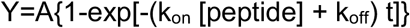

Where Y is the receptor binding signal and A is a constant. The dissociation constant K_d_ was calculated as k_off_/k_on_.

For steady-state analysis, the data were also fit to a one-site binding equation described in Roehrl *et al*., using Python for comparison (Roehrl et al., 2004).

### Crystallography

BAK:dF3 crystals were obtained by mixing 250 µM of BAK ΔN22 ΔC25 C166S in TBS buffer pH 7.5 with 250 µM of peptide (dissolved in water) in a 30 µl volume. Hanging drop crystals were grown at 4°C in polyethylene glycol (PEG) 3350 (20% w/v) and calcium acetate 0.2 M. BAK:dF2 crystals were obtained by mixing 250 µM of BAK ΔN22 ΔC25 C166S in TBS buffer pH 7.5 with 250 µM of peptide (dissolved in water) in a 30 µl volume. Hanging drop crystals were grown at 25 °C in 1.2 M disodium malonate, 0.1 M Tris, pH 8.5. BAK:dM2 crystals were obtained by mixing 250 µM of BAK ΔN22 ΔC25 C166S in TBS buffer pH 7.5 with 250 µM of peptide (dissolved in water) in a 30 µl volume. Hanging drop crystals were grown at 25°C in 3.5 M sodium formate. BAK:dF3, BAK:dF2, and BAK:dM2 crystals were frozen in the well solution and X-ray data were collected at Argon National Laboratories. Images were processed with HKL2000 or XDS and the structure was solved by molecular replacement with PHASER, searching for BAK monomer (PDB: 5VX0) without peptide. The final model was produced by rounds of building in COOT and refinement using PHENIX. PDB IDs will be updated.

### CD measurements

Peptide helicity was measured using an JASCO Circular Dichroism Spectrometer at 25 °C (unless indicated otherwise in **Table S6**. A single scan with speed of 50 nm/min at 0.5 nm increments (190 nm to 250 nm wavelengths) was recorded. The baseline signal for a buffer control (no peptide) was subtracted. Peptide samples were prepared in 10 mM sodium phosphate pH 7.0, with a final peptide concentration of 15 μM. The unfolded peptide concentration was determined by UV absorption in 6.0 M guanidine hydrochloride aqueous solution. As in (Shepherd et al., 2005) ellipticity [θ] was calculated as the mean residue ellipticity (MRE) in the units of deg cm^2^(10*dmol residue)^-1^ according to this equation:

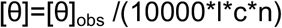

where [θ]_obs_ is the measured ellipticity in millidegrees [mdeg], **c** is the peptide concentration [mol L-1]), **n** is the number of residues, and **l** is path length (0.1 cm). The fractional helical content was calculated from the MRE at 222 nm and the number of backbone amides using the equation as in (Araghi et al., 2016):

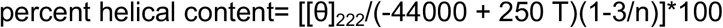

where n is the number of amino-acid residues in the peptide and T is the temperature in degrees Celsius.

### Structure-based design of BH3-only binders of BAK

We used the structure-based computational method dTERMen to design peptide binders of BAK (Zhou et al., 2020), following the approach of Frappier *et al*. (Frappier et al., 2019). In this work, we used version 35 of the dTERMen energy function and the same database of known structures as Zhou et. al (Zhou et al., 2020). To design BK3, we used BAK bound to Bim-h3Glg (PDB: 5VX0) as a template; Bim-h3Glg contains a non-native amino acid. Prior to peptide sequence design, we regularized the backbone of the peptide using the protocol TERMify, to improve compatibility with native amino-acid sequences. TERMify makes small adjustments to the backbone of a given structure to maximize similarity to common structural motifs in known proteins. A single cycle of the protocol consists of the following steps. First, the backbone structure is divided into overlapping fragments. Each fragment is then searched against a database of known structures to identify the top *N* matches, ranked by lowest RMSD. Finally, the method Fuser is used to update the coordinates of the original backbone to maximize similarity to the structural matches (Swanson et al., 2022). This process can be repeated to introduce progressively larger changes to the backbone, but in practice we have found that after ∼50 cycles only minor structural changes are observed between cycles. We defined single-residue and residue-pair fragments from the structure at the beginning of every cycle. For each residue in the peptide, *r*_*i*_, we defined a self-fragment consisting of *r*_*i*_ as well as flanking residues *r*_*i*−1_ and *r*_*i*+1_. For each residue *r*_*j*_ with the potential to contact *r*_*i*_, we defined a pair fragment consisting of the contacting residues and the residues flanking them (*r*_*i*−1_, *r*_*i*_, *r*_*i*+1_, *r*_*j*−1_, *r*_*j*_, *r*_*j*+1_). Residues were considered to have the potential to make a contact if their contact degree was greater than 0.01 (Holland et al., 2018). We searched a database of 12,657 non-redundant known structures for matches to the fragments using FASST, which guarantees that all structural matches within a given RMSD cutoff are identified and returned (Zhou & Grigoryan, 2015). We used a size and topology dependent function to define the RMSD cutoff used in structural searches, as in Zhou et al. (Zhou et al., 2020). We modified TERMify to account for the fixed structure and amino-acid sequence of BAK, so that only the peptide backbone geometry was updated. Single-residue fragments were defined from peptide residues only, and pair fragments were defined only for peptide-peptide and peptide-protein potential contacts. We used the sequence of BAK as an additional constraint when searching for structural matches. Specifically, for a pair fragment involving a peptide residue *r*_*i*_ and protein residue *r*_*j*_, we required that the residue corresponding to *r*_*j*_ in the match have the same amino acid as BAK. As in Swanson et. al, we fixed the structure of BAK when applying Fuser, so that only the structure of the peptide backbone was updated. When using TERMify to relax the structure of the peptide in PDB: 5VX0, we searched for 10 matches to each fragment in each of 100 cycles. The code detailing the TERMify procedure is provided at https://github.com/Grigoryanlab/Mosaist.

### dTERMen scoring of human proteome sequences

BH3-containg sequences from the human proteome were scored using solved crystals structures of BAK:peptide complexes and dTERMen version 35. Because energy scores are not directly comparable between structures, we first ranked sequences according to their energy scores generated with the same structure and then compared rank order among all three structures to obtain an overall rank position.

### Cavity detection

Cavities were detected and measured using F-pocket (Guilloux et al., 2009) available at: https://github.com/Discngine/fpocket.

### siRNA transfection

24 hours before siRNA transfection, HeLa cells were seeded in 6-well plates and incubated overnight at 37°C. Cells were transfected with siRNA (final amount per well 25pmol) using Lipofectamine RNAiMAX (ThermoFisher 13778075) according to manufacturer’s instructions. Cells were collected 72 hours later for BH3 profiling and western blotting.

BAX siRNA: ThermoFisher Silencer® Select, siRNA ID s1888

BAK siRNA: ThermoFisher Silencer® Select, siRNA ID s1881

Scramble siRNA: Dharmacon Horizon Discovery, ON-TARGETplus Non-targeting Control Pool (D-001810-10-05)

### BH3 profiling

For each siRNA transfection sample, 2 million cells were centrifuged at 400g for 5 minutes and subjected to BH3 profiling as previously described (Fraser et al., 2019). In short, BH3 peptides in mannitol experimental buffer (MEB) (10 mM HEPES pH 7.5, 150 mM mannitol, 50 mM KCl, 0.02 mM EGTA, 0.02 mM EDTA, 0.1% BSA, 5 mM succinate) with digitonin were deposited into each well in a 96 well plate. Single cells were resuspended in MEB and added to each treatment well and incubated for 60 minutes at 28°C. Peptide exposure was terminated with formaldehyde and cells were stained overnight with AF647 conjugated cytochrome c antibody (BioLegend 612310) and DAPI. Cytochrome c positivity was measured on ThermoFisher Scientifice Attune NxT Flow Cytometer.

### Immunoblotting

Protein lysates were obtained by cell lysis in RIPA buffer (Boston BioProducts 115) with protease inhibitor (Roche 11697498001) and phosphatase inhibitor (ThermoFisher A32957). Protein loading was measured by BCA Protein Assay (ThermoFisher 23227). Protein samples were electrophoretically separated in precast gels (BioRad Mini-PROTEAN TGX Gels). Protein was transferred to PVDF membrane using Bio-Rad Trans-blot Semi-Dry transfer cell, blocked with 5% milk, and incubated overnight with primary antibody (BAX: Cell Signaling 2772S, BAK: Millipore Sigma 06-536, GAPDH: Cell Signaling 2118L). Secondary antibody (Peroxidase-linked anti-Rabbit: Cytiva Lifescience NA934) incubation and Pico development (ThermoFisher 34579) was performed before imaging on (Invitrogen iBright FL1500 Imaging System).

## Acknowledgements and Funding

This work was funded by the National Institute of General Medical Sciences (NIGMS) award R01 GM110048 to A.E.K., MIT School of Science Fellowship in Cancer Research award to F.A., and the John W. Jarve (1978) Seed Fund for Science Innovation (MIT) award to F.A. and A.E.K. Part of this work is based upon research conducted at the Northeastern Collaborative Access Team beamlines, which are funded by the NIGMS (P30 GM124165). The Eiger 16M detector on 24-ID-E is funded by NIH-ORIP HEI grant S10OD021527. This research used resources of the Advanced Photon Source, a U.S. Department of Energy (DOE) Office of Science User Facility operated for the DOE Office of Science by Argonne National Laboratory under Contract No. DE-AC02-06CH11357. This work was supported in part by the Koch Institute Support (core) Grant P30-CA14051 from the National Cancer Institute. We thank the Koch Institute’s Robert A. Swanson (1969) Biotechnology Center for technical support, specifically the Biopolymers and Proteomics Facility and the Flow Cytometry Core Facility. We also thank the MIT Structural Biology Core Facility, under the leadership of R.A.G., and we thank M. Carney for her assistance with binding assays.

## Author Contributions

F.A. and A.E.K. conceptualized the project, designed and analyzed experiments, and wrote the paper. F.A. performed the biochemical and structural experiments. S.Y. conducted cell-based assays. R.A.G. directed and assisted with crystallography experiments and analysis. S.S. conducted peptide computational design and RMSD calculations. D.G. assisted with cell-surface display experiments. B.G.S. assisted with crystallography and assay development. K.A.S. helped with the design and interpretation of cell-based experiments.

