## Supplementary Information for "Discovery of diverse human BH3-only and non-native peptide binders of pro-apoptotic BAK indicate that activators and inhibitors use a similar binding mode and are not distinguished by binding affinity or kinetics"

**A)**

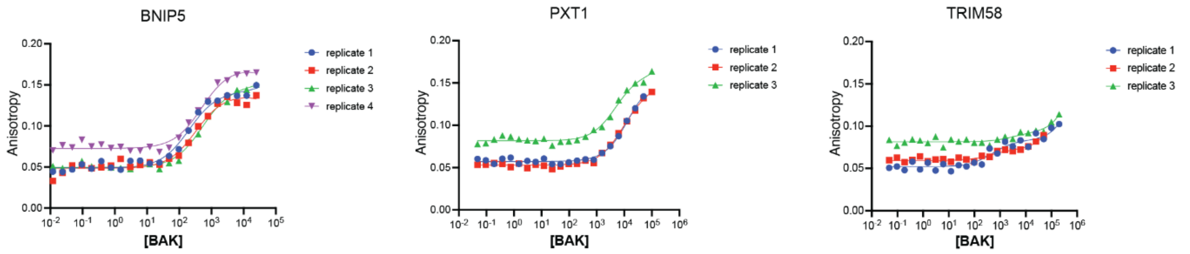


| Peptide | Replicate #1 | Replicate #2 | Replicate #3 | Replicate #4 | Average $K_d$ |
|---|---|---|---|---|---|
| BNIP5 | 0.2 µM | 0.3 µM | 0.6 µM | 0.4 µM | 0.4 µM +/- 0.2 |
| PXT1 | 14 µM | 12 µM | 7 µM | - | 11 µM +/- 4 |

**B)**

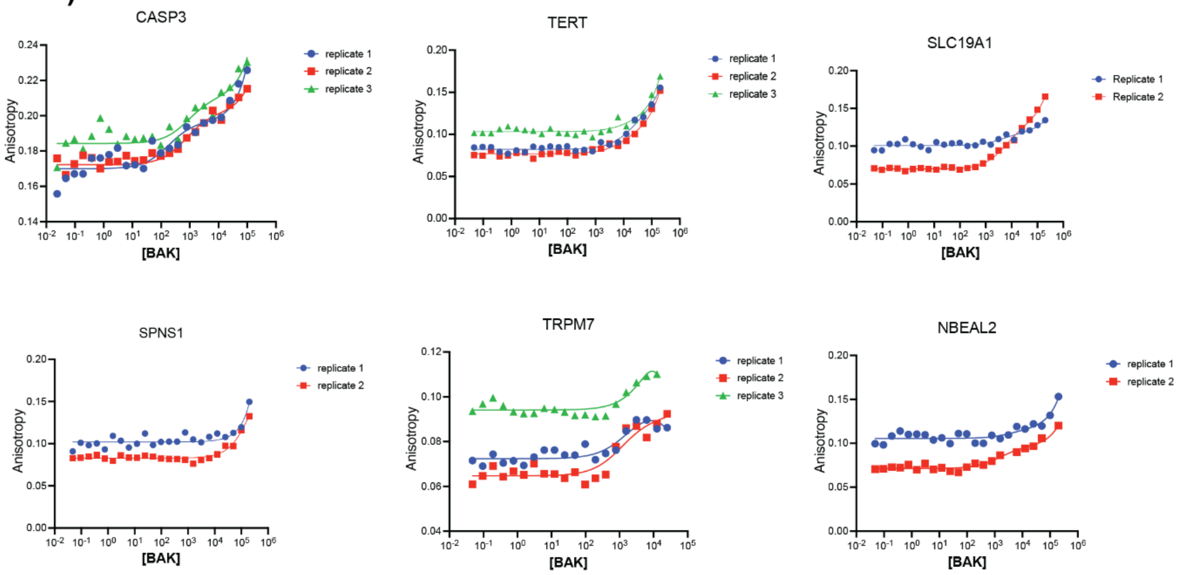


**Figure S1. Candidate BH3-only peptides bind BAK with a range of affinities**. Peptides that bind anti-apoptotic proteins MCL-1, BCL-$x_L$, BCL-2, BCL-W, and BFL-1 were tested for binding to recombinant human BAK using fluorescence polarization. **A)** BNIP5 and PXT1 showed the tightest binding with dissociation constants of 400 nM and 11 µM, respectively. **B)** CASP3, TERT, and SLC19A1 showed weak binding up to 200 µM BAK. SPNS1, TRPM7, and NBEAL2 were the weakest binders with unmeasurable affinities. SNTG2, TXDC11, POFUT2, DDX4, and MINA did not bind BAK (data not shown).

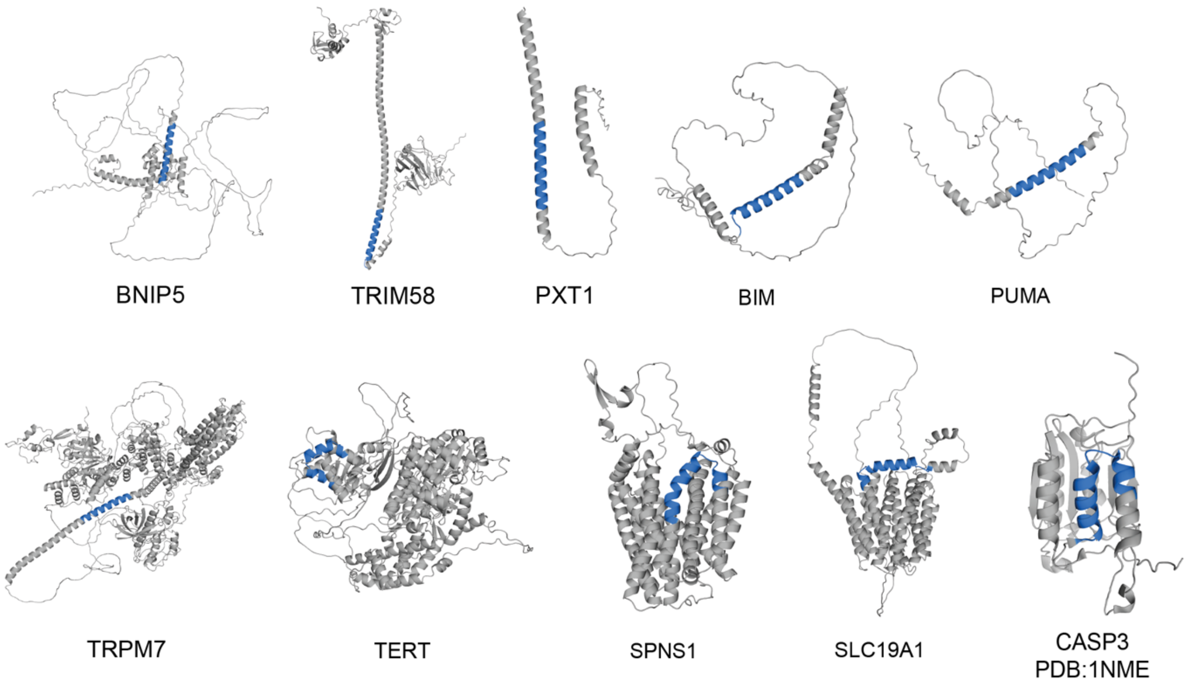


**Figure S2. AlphaFold structure predictions for human BAK-binding proteins**. The candidate BH3 motif is highlighted in blue. NBEAL2 in progress.

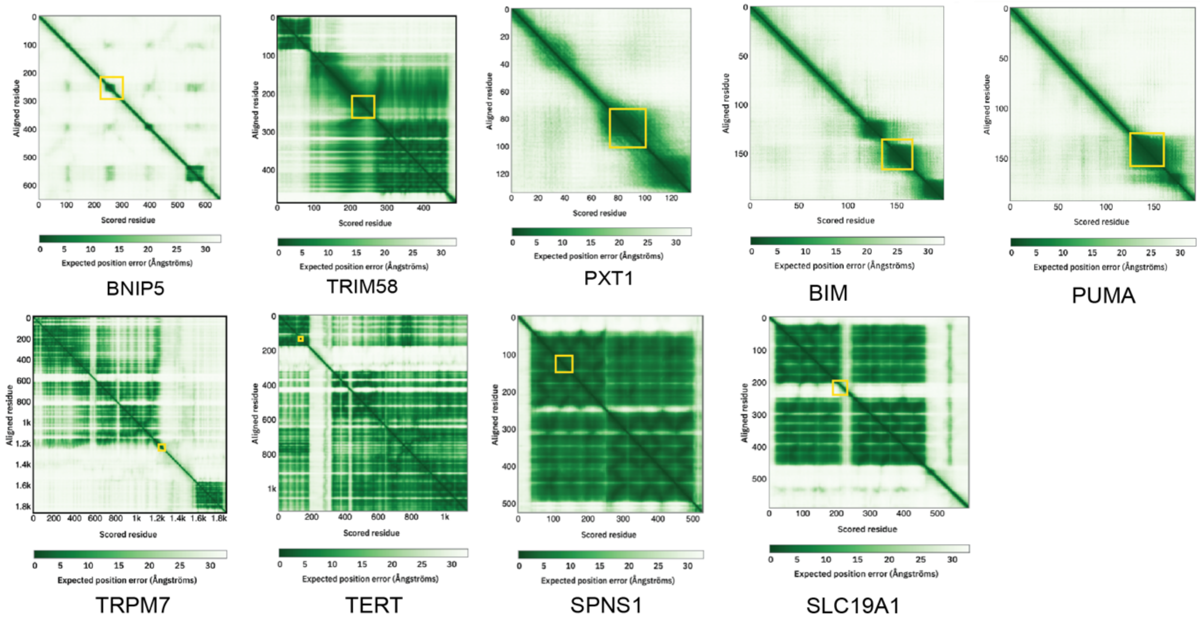


**Figure S3. AlphaFold Predicted Aligned Error for predicted structures of human proteins.** The candidate BH3 motif is indicated with a yellow box. NBEAL2 in progress.

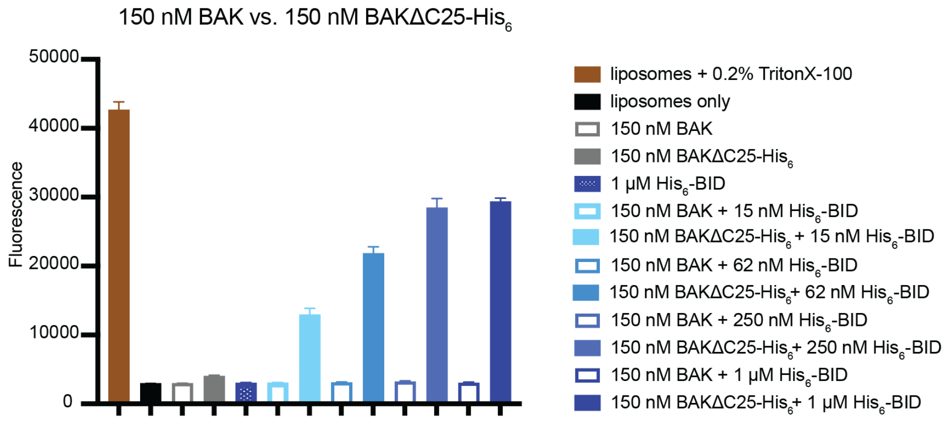


**Figure S4. Localization of BAK to the liposome membrane is necessary for BID BH3-triggered dye release.** Liposomes containing ANTS/DPX dye and 18:1 DGS-NTA ($Ni^{2+}$) lipid were incubated with either 150 nM of BAK ΔN22 ΔC25 C166S (indicated as BAK) or 150 nM of BAKΔC25-$His_6$ and increasing concentrations of $His_6$-SUMO-BID BH3 (indicated as $His_6$-BID). Whereas BAKΔC25-$His_6$ showed BID concentration-dependent dye release, BAK without a $His_6$ tag did not lead to dye release even when treated with 1 μM $His_6$-BID. 0.2% Triton X-100 detergent (brown) was included as a positive control for dye release. Plots show the fluorescence signal at 1 hour, for experiments run in triplicate, with error bars representing standard deviations.

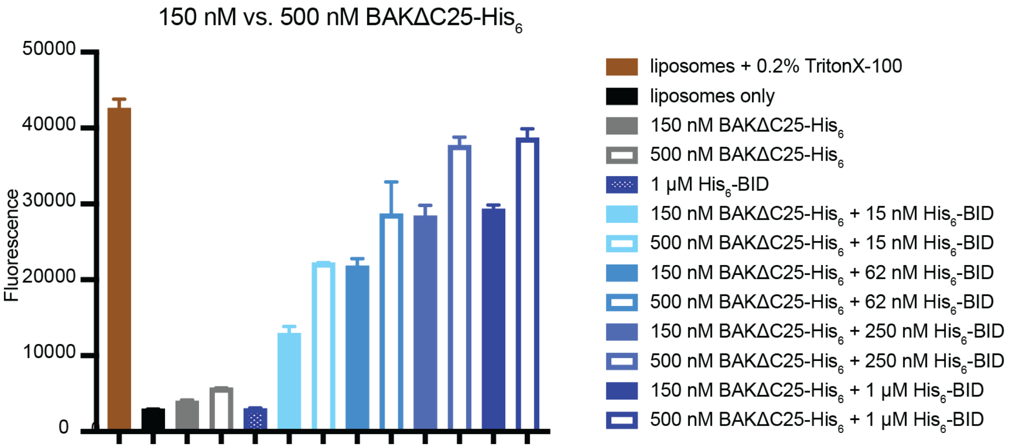


**Figure S5. Dye release increases with increasing concentrations of BAK.** For all tested concentrations of His-SUMO-BID BH3 (indicated as $His_6$-BID), greater dye release was observed for 500 nM vs. 150 nM BAKΔC25-$His_6$. No dye release was observed for BAKΔC25-$His_6$ in the absence of $His_6$-BID at either 150 or 500 nM. Plots show the fluorescence signal at 1 hour for experiments run in triplicate, with error bars representing standard deviations.

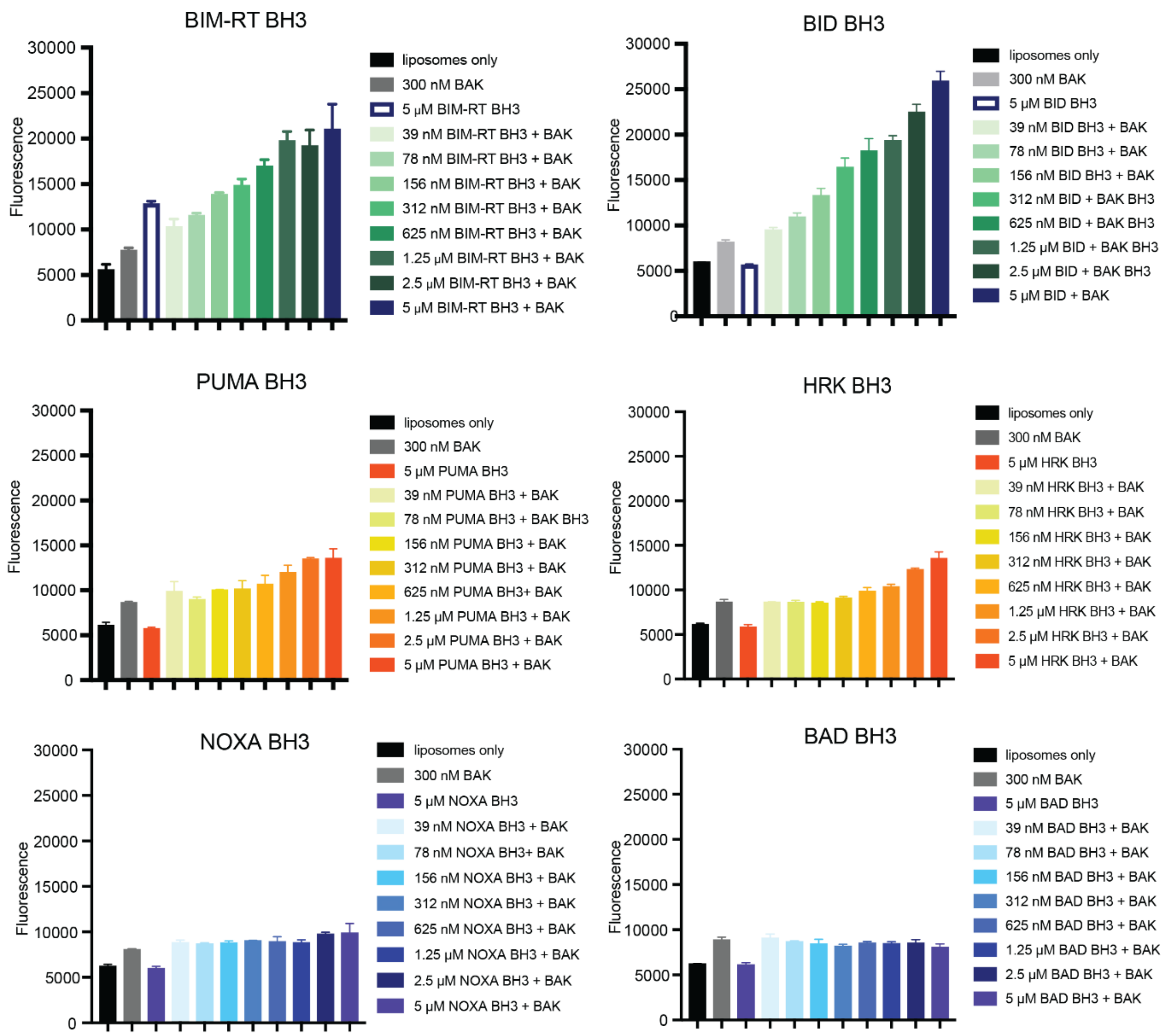


**Figure S6. Peptides from BCL-2 BH3-only proteins show differences in BAK activation function.** BH3 peptides BID, BIM, PUMA, HRK, Y-NOXA, and BAD were tested for activation. 300 nM BAKΔC25-$His_6$ (indicated as BAK in the figure) was incubated with BH3 peptides at concentrations ranging from 39 nM to 5 μM. BH3 peptides were grouped into activators (green), weak activators (orange) and non-activators (blue). Plots show the fluorescence signal at 1.5 hours for experiments run in triplicate, with error bars representing standard deviations. Peptide sequences are in **Table S5.**

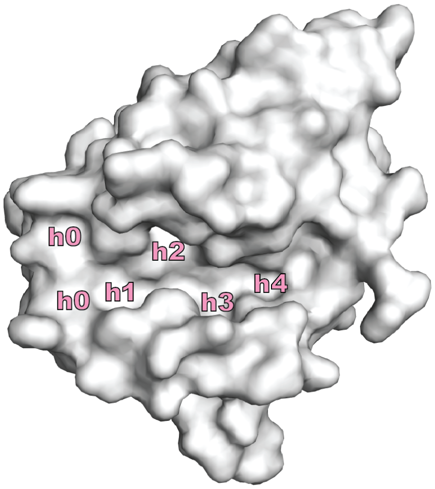


**Figure S7. BAK contains 5 hydrophobic pockets referred to as h0, h1, h2, h3, and h4.** Surface representation of BAK (PDB:5VX0) with hydrophobic pockets in the canonical peptide binding groove labeled in pink.

| | Sequence |
|---|---|
| | 1 \| 2 \| 3 \| 4 \| 5 |
| | cdefgabcdefgabcdefgabcdefgabc |
| | h0 h1 h2 h3 h4 |
| PUMA | EEQWAREIGAQLRRMADDLNAQYERR |
| PUMA M3dI | EEQWAREIGAQLRRIADDLNAQYERRR |
| NOXA | PAELEVECATQLRRFGDKLNFRQKLL |
| NOXA C2dI | YPAELEVEIATQLRRFGDKLNFRQKLL |

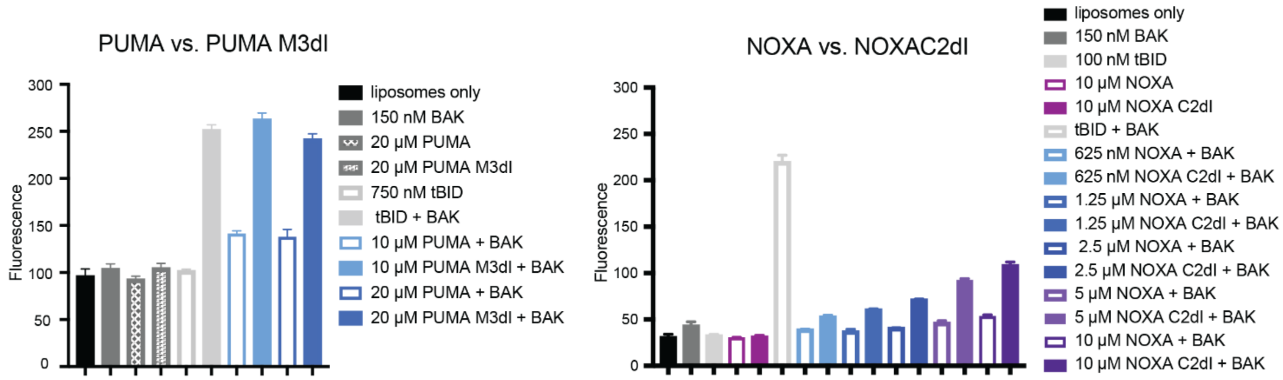


**Figure S8. Point mutations at hydrophobic positions increase the activation potency of BH3 peptides from proteins NOXA and PUMA.** Using heptad notation, peptide residues 1g and 2a, 2d, 3a, 3d and 4a bind to hydrophobic pockets h0, h1, h2, h3, and h4, respectively. Substitution of Met to Ile in Puma BH3 peptide (PUMA M3dI) and Cys to Ile in NOXA BH3 peptide (NOXA C2dI) increased activation potency. Dye containing liposomes were incubated with 150 nM BAKΔC25-His$_6$ (indicated as BAK) and peptides with concentrations ranging from 625 nM to 10 μM. Plots show the fluorescence signal at 1 hour for experiments run in triplicate, with error bars representing standard deviations.

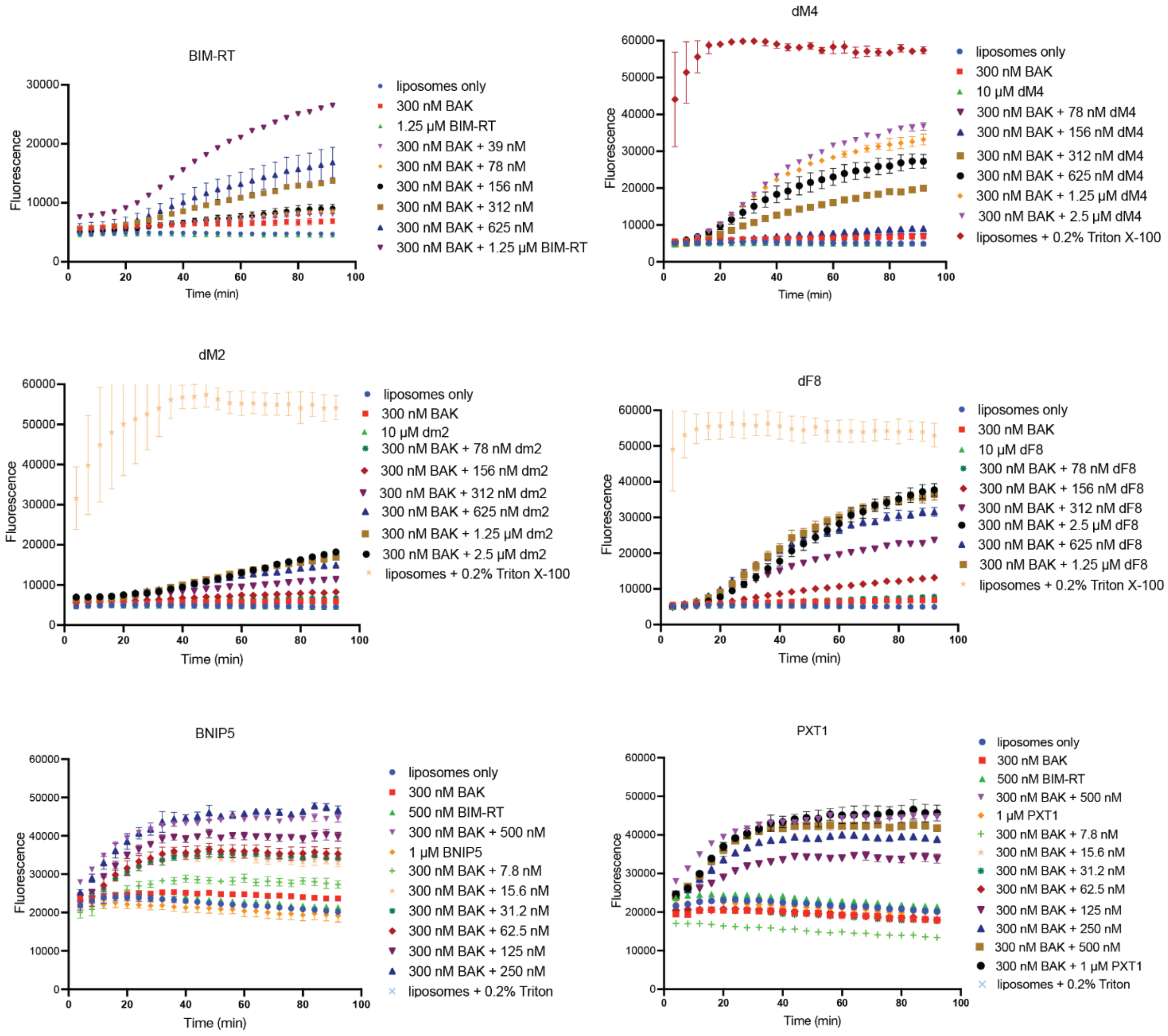


**Figure S9. Raw liposome data of BAK activator peptides.** Data correspond to **Figure 3** in the main text.

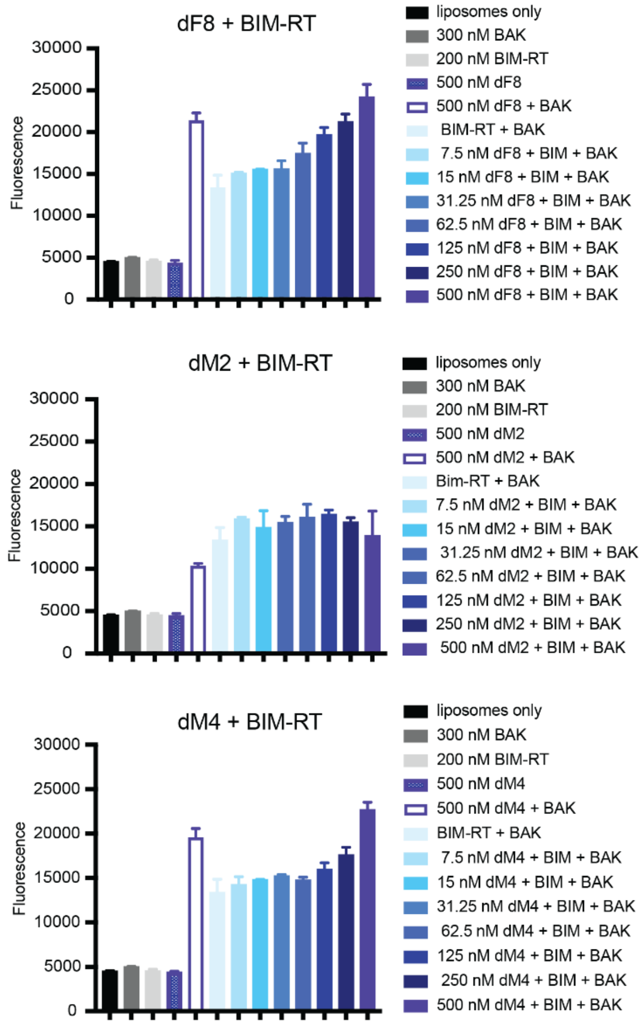


**Figure S10. Peptides dF8, dM2, and dM4 act as activators alone and in the presence of BIM-RT BH3.** Liposomes containing dye were incubated with 300 nM BAKΔC25-His$_6$ (indicated as BAK in the figure), 200 nM BIM-RT (activator), and 7.5 – 500 nM dF8, dM2, or dM4 peptides. Plots show the fluorescence signal at 2 hours. All peptide combinations displayed greater fluorescence compared to 300 nM BAKΔC25-His6 and either 200 nM BIM-RT or 500 nM dF8, dM2, or dM4 peptide alone.

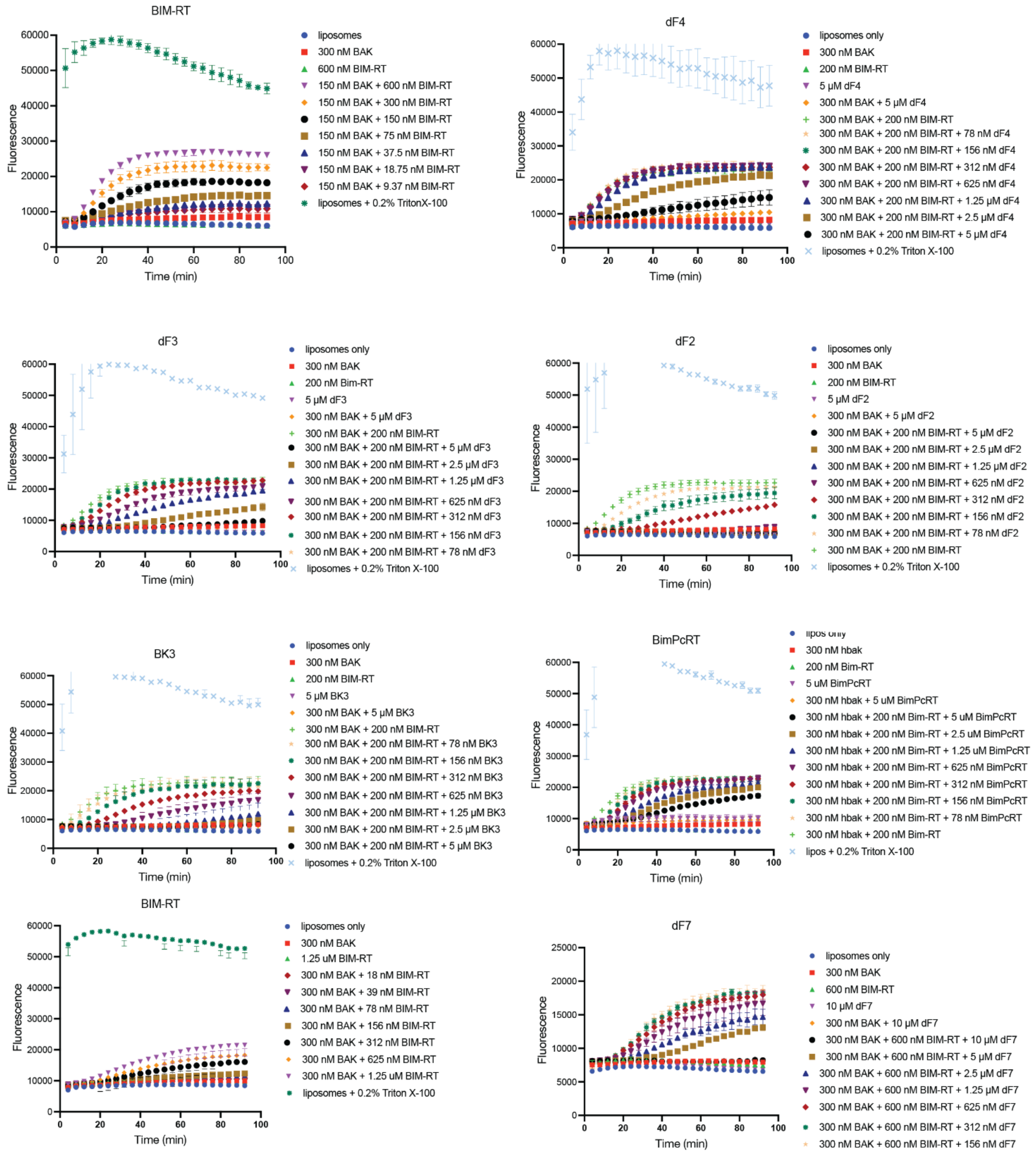


**Figure S11. Raw liposome data for BAK inhibitor peptides.** Data correspond to **Figure 4** in the main text.

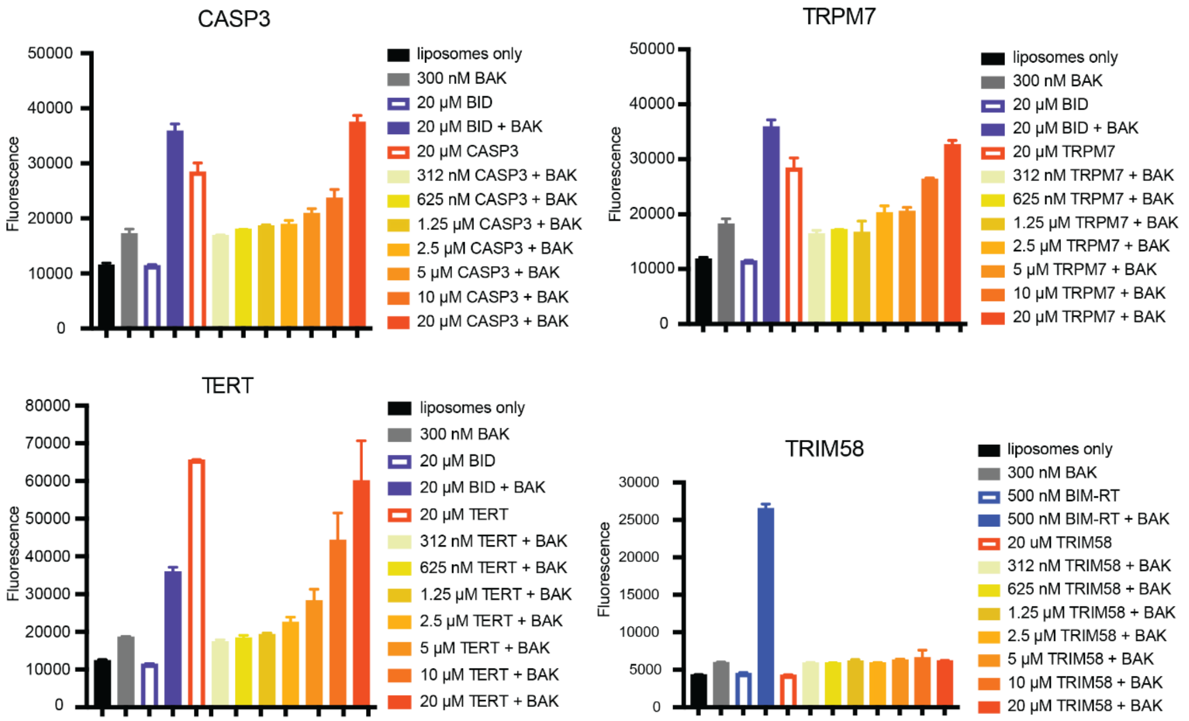


**Figure S12. CASP3, TERT, and TRPM7 BH3 peptides show BAK-independent membrane disruption at high concentrations.** BID-Y 25mer is a positive control showing concentration dependent activation in the presence of 300 nM BAKΔC25-His$_6$ (indicated as BAK in figure). CASP3, TERT, and TRPM7 peptides show membrane disruption at concentrations greater than 5 μM, 2.5 μM, and 5 μM, respectively. TRIM58 does not show activation at 20 μM.

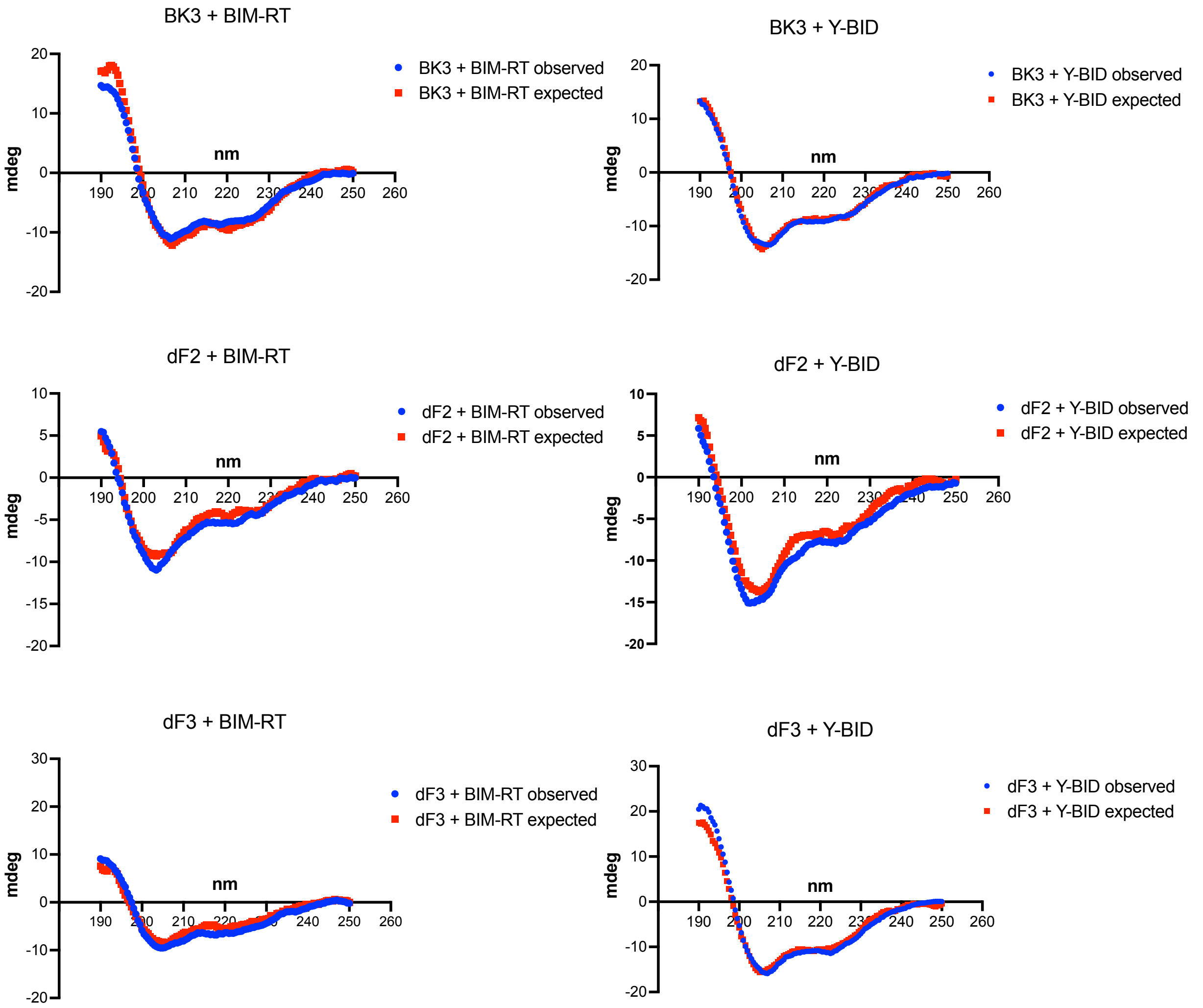


**Figure S13. Circular dichroism (CD) experiments do not support hetero-dimerization of inhibitor peptides with BIM-RT.** CD spectra were collected for each peptide at a concentration of 15 μM. Summing the CD signals for the indicated peptides gave the expected spectra, which are plotted in red. For comparison with the signal expected from a physical mixture of two non-interacting peptides, 15 μM BIM-RT or Y-BID was mixed with 15 μM BK3, dF2, or dF3 to obtain the observed CD spectra, which are plotted in blue. The observed spectra (blue) showed minimal differences from the spectra expected for non-interacting peptides (red).

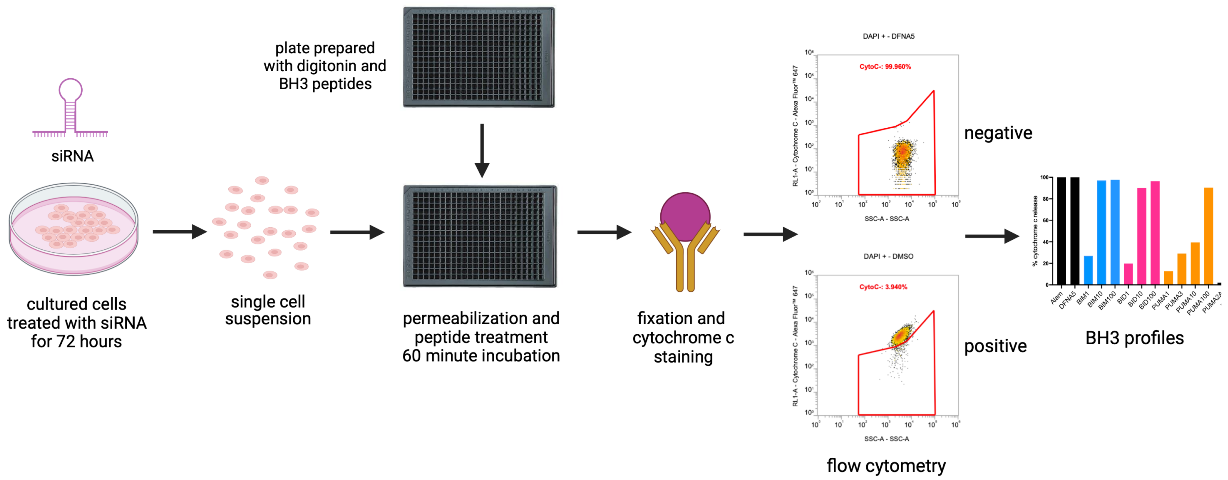


**Figure S14. Diagram of BH3 profiling assay.**

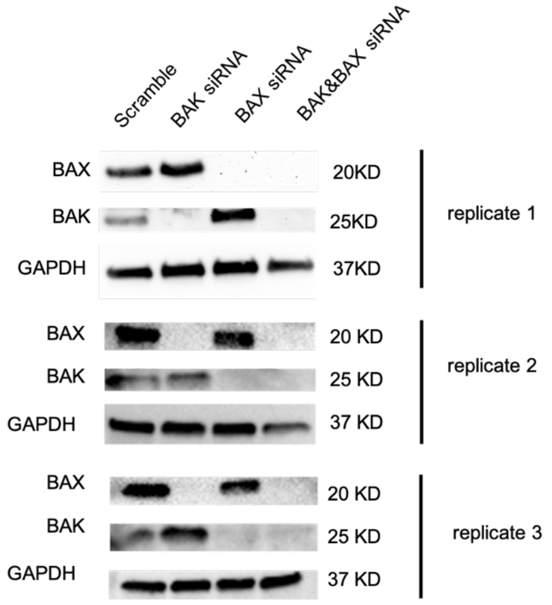


**Figure S15. Western blots performed after 72-hour siRNA treatment of HeLa cells show levels of BAK and BAX, with GAPDH as a loading control.**

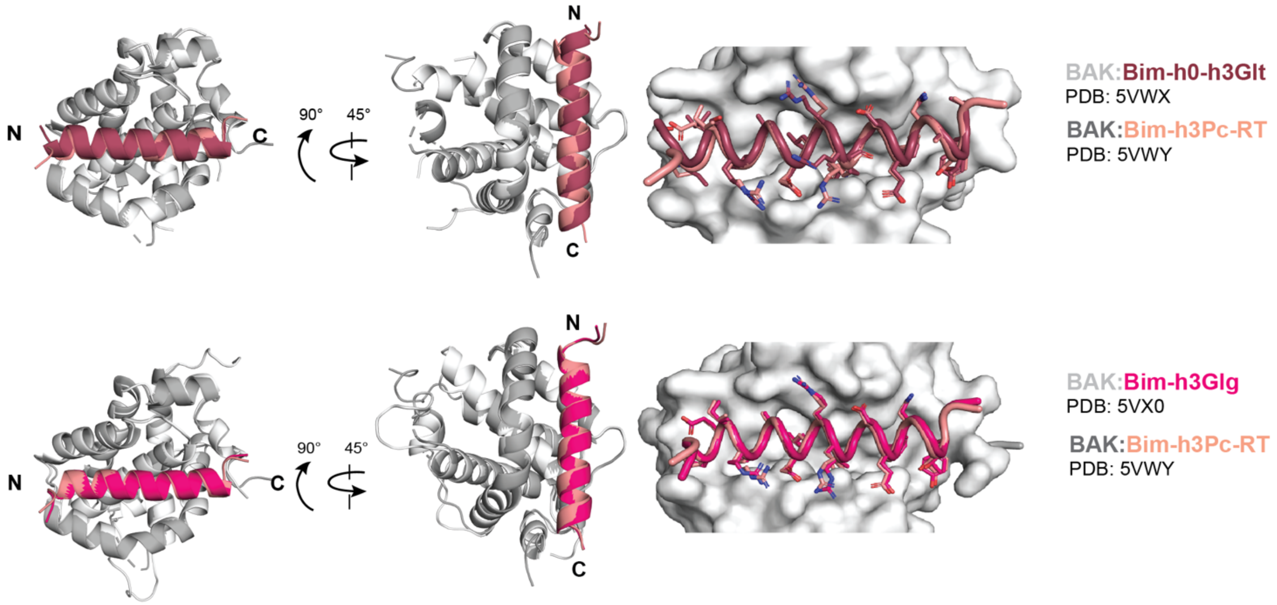


**Figure S16. Peptide inhibitors containing non-natural amino acids bind to BAK in very similar binding modes.** Structures are from (Brouwer et al., 2017). The inhibitor peptide sequences are the same, except for the non-natural amino acid at the 3d position that binds in the h3 pocket. Top) Comparison of inhibitors bound to domain-swapped core-latch BAK dimer structures: Bim-h0-h3Glt (raspberry, PDB: 5VWX) and Bim-h3Pc-RT (salmon, PDB: 5VWY). Bottom) Comparison of inhibitors bound to monomeric BAK (PDB:5VX0) and core-latch dimer BAK (PDB: 5VWY). Bim-h3Glg is in hot pink and Bim-h3Pc-RT is in salmon.

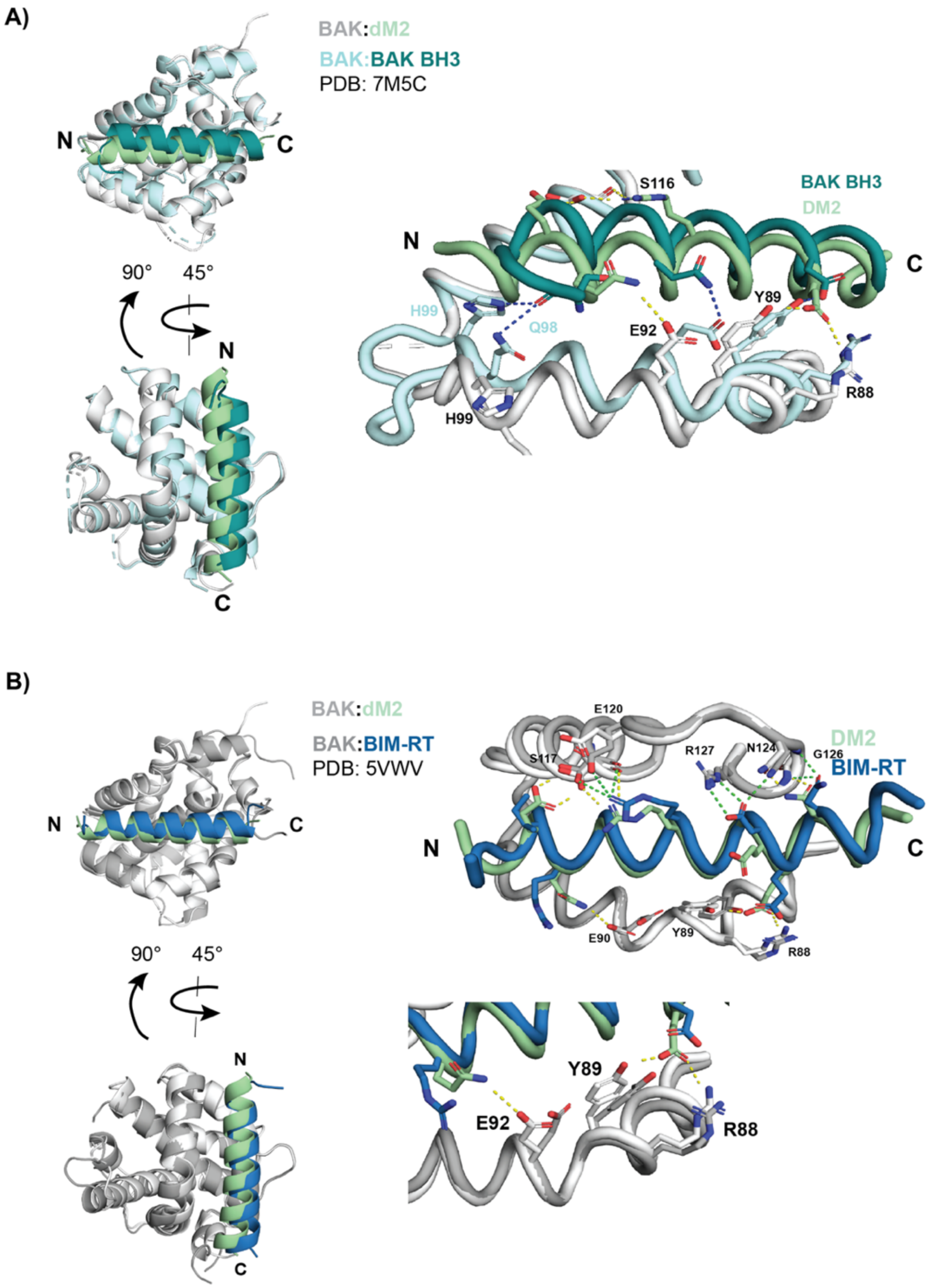


**Figure S17. Activator dM2 binds differently compared to BAK BH3 and BIM-RT activator peptides. A)** Comparison of dM2 (pale green) with BAK BH3 (deep teal) shows a shift of the N-terminus of the BAK BH3 peptide and the BAK $\alpha 3$ helix that forms part of the binding site. This difference in peptide binding mode is accompanied by a difference in polar contacts (yellow for DM2 and navy blue for BIM-RT). Specifically, BAK BH3 peptide forms contacts with H99 and Q98 on BAK that are not made by dM2. **B)** Superimposed crystal structures of dM2 (pale green) and BIM-RT (sky blue) show slightly different positioning of the C-termini of the peptides. This difference allows dM2 to make polar contacts with residues E92, Y89, and R88 on BAK.

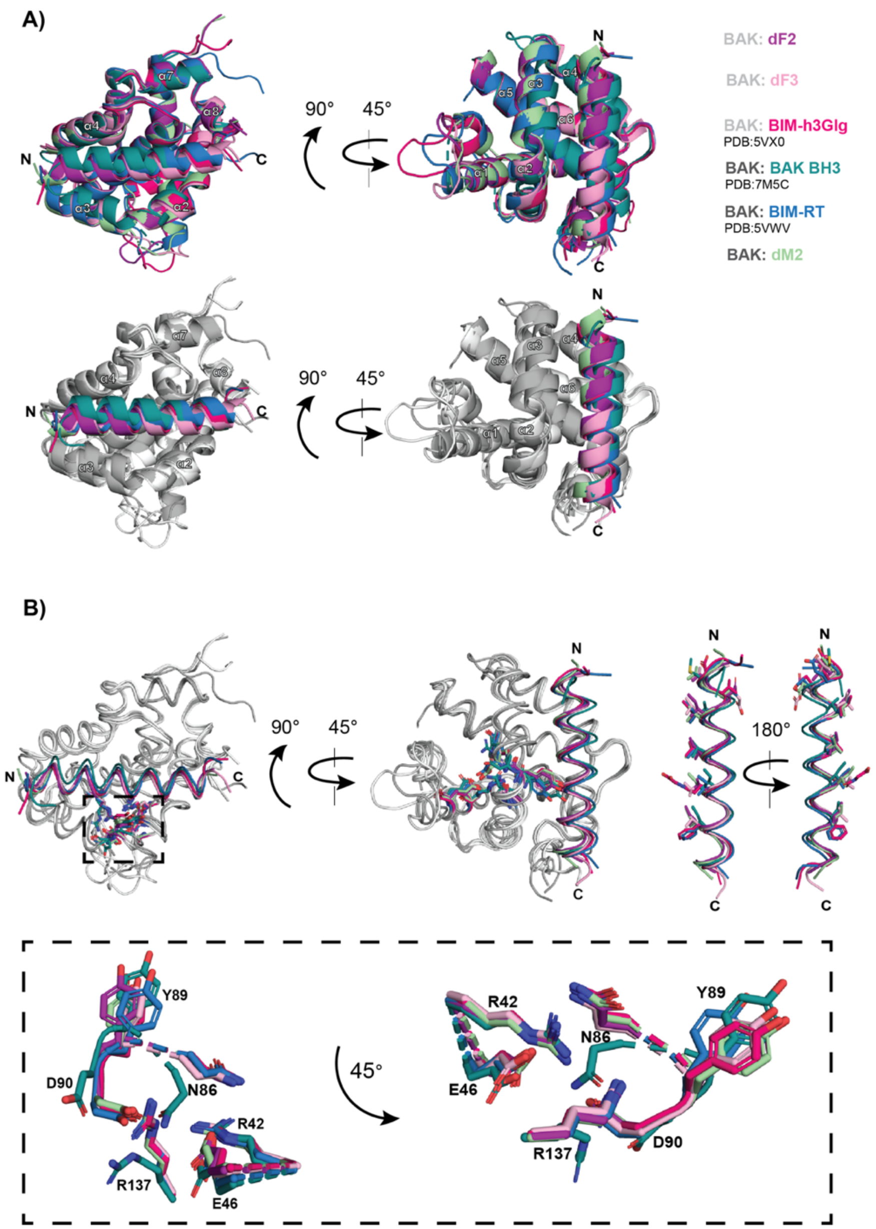


**Figure S18. Activators and inhibitors of BAK bind with no systematic differences in structure. A)** Superposition of complexes of BAK bound to activators (BAK BH3, BIM-RT, and dM2 peptides) and inhibitors (dF2, dF3, and BIM-h3Glg peptides). BAK BH3 (deep teal, PDB: 7M5C) shows the greatest deviation compared to the rest of the structures, with a shift of the N terminus of the peptide and the $\alpha$3 helix of BAK. (Top) BAK and corresponding peptide shown in the same color. (Bottom) Same figure as in top, but with BAK colored in shades of grey for clearer visualization. **B)** (Left) Ribbon representation of cartoon in A). Residues involved in a previously reported electrostatic network are depicted with sticks and enclosed in a dotted rectangle. A closer look at the electrostatic network involving residues N86, Y89, D90, R42, E46, and R137 viewed from two different perspectives differing by 45°. (Right) All six peptides are superimposed with hydrophobic residues depicted with sticks.

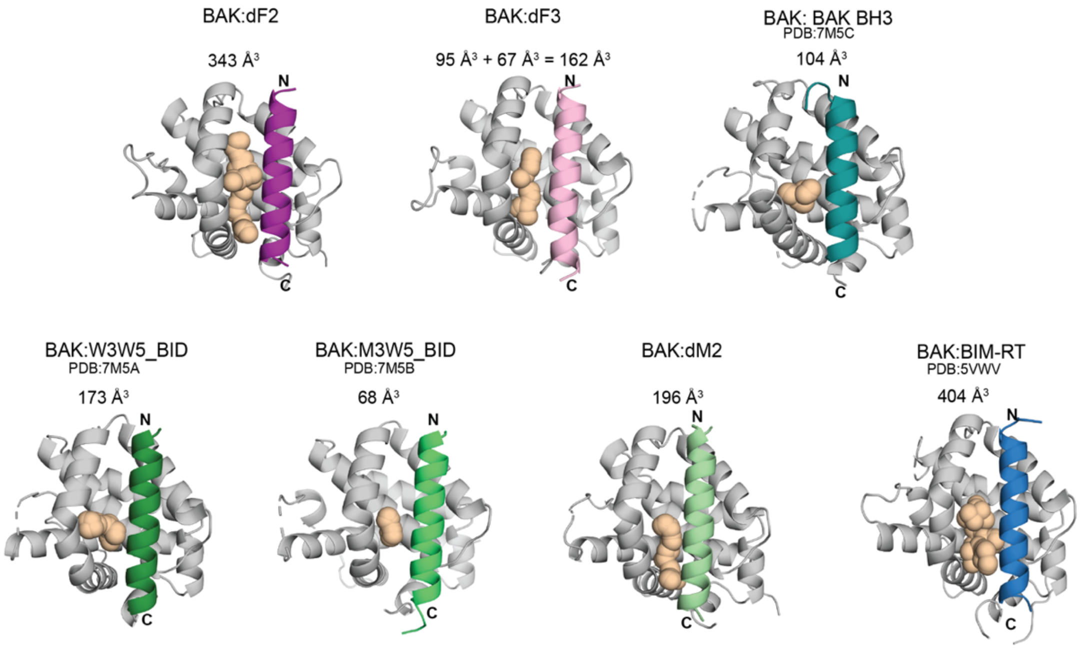


**Figure S19. Cavity sizes in BAK:peptide complexes do not correlate with function.** The program F-pocket was used to detect and quantify cavity volumes (indicated with wheat-colored spheres) using a minimum probe radius of 3.4 Å and a maximum probe radius of 6.2 Å. Inhibitor peptides are shown in purple and pink, and activator peptides are shown in shades of green and blue.

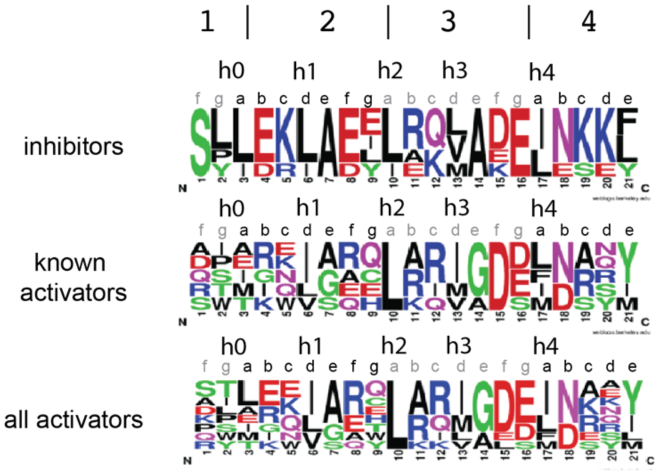


**Figure S20. Sequence logos of activators and inhibitors show few residue preferences.** Peptide sequence logos were generated for inhibitors (dF2, dF3, BK3, dF4, dF7), known activators (BID, BIM, PUMA, BAK, and BAX BH3 regions), and all activators (BID, BIM, PUMA, BAK BH3, BAX BH3, dM2, dF8, and dM4).

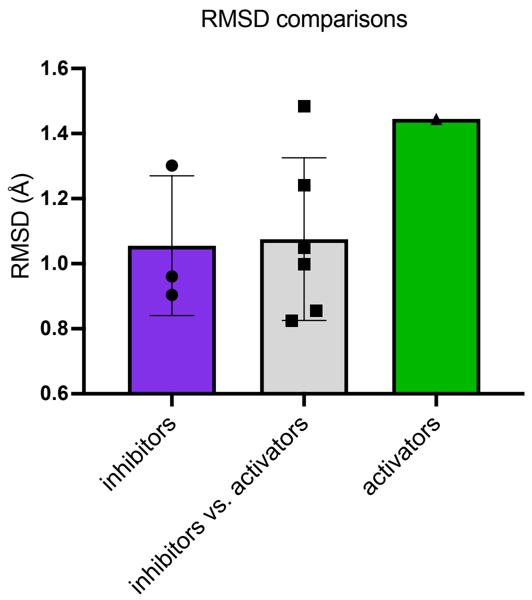


**Figure S21. Pairwise RMSD calculations of activator vs. inhibitor peptides show an average RMSD of 1.1 Å.** Crystal structures of BAK:peptide complexes were aligned based on BAK and then differences between peptide binding geometries were compared based on the all-backbone-atom RMSD values for inhibitors vs. inhibitors (purple), inhibitors vs. activators (grey), or activators vs. activators (green). Structures used for analysis included (BAK:dF2, BAK:dF3, BAK:Bim-h3Glg - PDB:5VX0, BAK:BIM-RT - PDB:5VWV, and BAK:dM2).

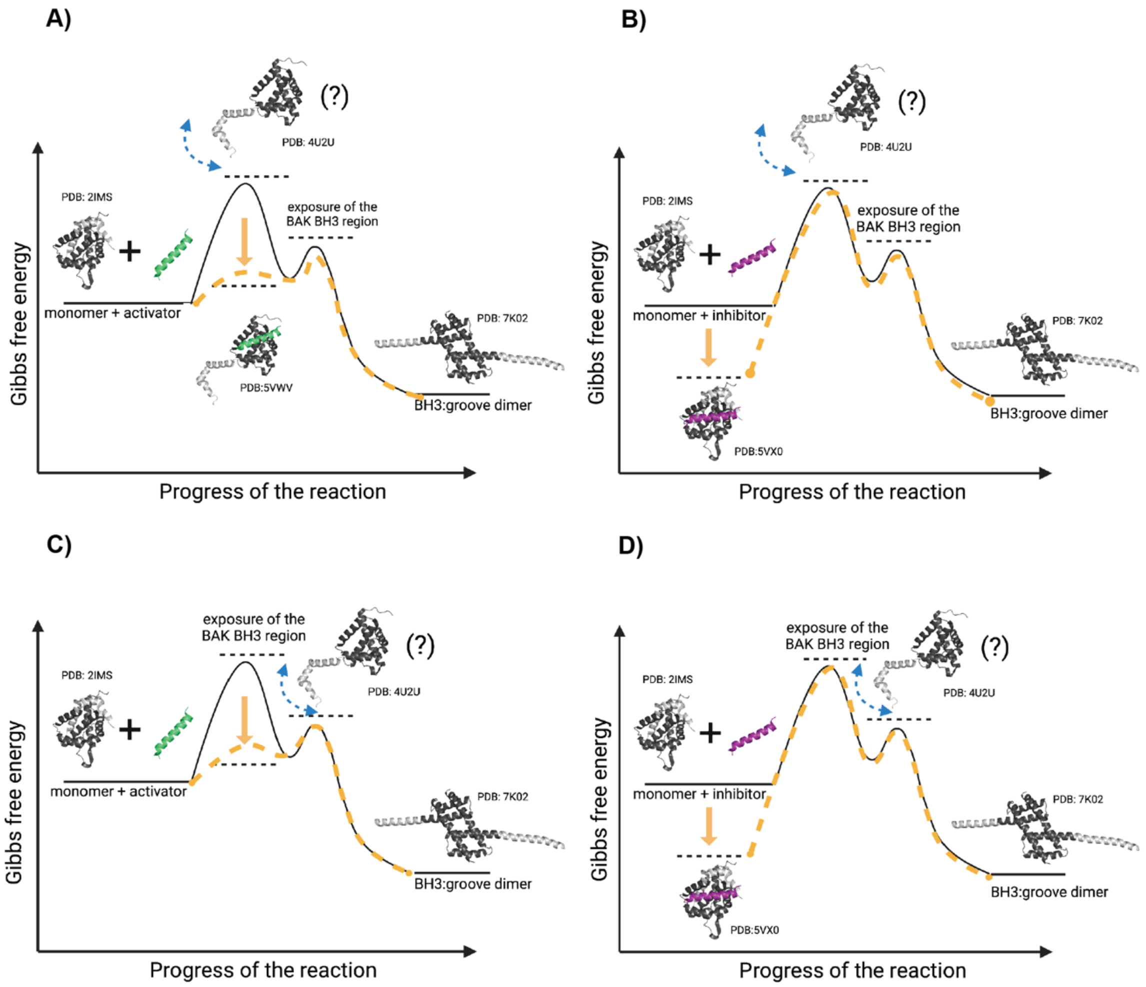


**Figure S22. Free energy diagram to explain differences between activators and inhibitors of BAK**. Depicted are putative intermediate crystal structures of BAK with the core in dark grey and the latch in light grey as described for **Figure 9**. Exposure of the BAK BH3 is depicted as occurring **top)** after release of the latch from the core and **bottom)** before release of the latch from the core.) **A)** and **C)** illustrate BAK activators stabilizing the transition state. **B)** and **D)** illustrate BAK inhibitors stabilizing the ground state.

### Tables

| PEPTIDE NAME | SEQUENCE | SOURCE | FACS PROFILE |
|---|---|---|---|
| | 1 \| 2 \| 3 \| 4 \| 5 | | |
| | cdefgabcdefgabcdefgabcdefgabcdefg | | |
| | h0 h1 h2 h3 h4 | | |
| dF4 | ---SLLEKLAEYLRQMADEINKKYVK------- | Frappier, V. *et al.* | high |
| dF3 | ---SLLEKLAEELRQLADELNKKFEK------- | Frappier, V. *et al.* | high |
| dF7 | ---SLLEKLAEELAQLADELNKKFEK------- | Frappier, V. *et al.* | high |
| BIM | -DMRPEIWIAQELRRIGDEFNAYYAR------- | BH3-only protein | high |
| dM2 | --APYLEQVARTLRKIGEEINEALR-------- | Frappier, V. *et al.* | high |
| dM3 | --DKTLEEIARELAKLAEEIDKEI--------- | Frappier, V. *et al.* | high |
| dF2 | ---SYIDKIADLIRKVAEEINSKLE-------- | Frappier, V. *et al.* | high |
| BK3 | --KSPLERLAEILEKVAKEIEKELGP------- | dTERMen (this work) | high |
| dM4 | --DKTLEEIARWLARLALEIDKEI--------- | Frappier, V. *et al.* | high |
| dF8 | ---SLLEKLAEYLAQMGDEINKKYVK------- | Frappier, V. *et al.* | medium |
| dF6 | ---SYIDKIADLIDKVVEEINSKLE-------- | Frappier, V. *et al.* | medium |
| dM5 | --APKEKEVARTLIKIGEEINEALK-------- | Frappier, V. *et al.* | medium |
| dM7 | --DKTLEEIARELLKLALEIDKEI--------- | Frappier, V. *et al.* | medium |
| dM1 | --APKEKEVAETLRKIGEEINEALK-------- | Frappier, V. *et al.* | low |
| dM9 | ---DIEQEIAEALKEVADELSKAIED------- | Frappier, V. *et al.* | low |
| dM6 | --APYLEQVARTLLHIGMEINEALR-------- | Frappier, V. *et al.* | low |
| dF5 | ---SYVDKIADLMKKVAEKINSDLT-------- | Frappier, V. *et al.* | low |
| MF2 | --GRWIDQIAQFLRRIGDHIEKYI--------- | Jenson, J.M. *et al.* | none |
| dM10 | ---DVVLSVAETLRELADRLYEEINT------- | Frappier, V. *et al.* | none |
| M1 | --GRSELEVVQELVRIGDIVVAYF--------- | Jenson, J.M. *et al.* | none |
| MF6 | --GRRVDEIAQILRRIGDNVTTYI--------- | Jenson, J.M. *et al.* | none |
| XF2 | --GRREVWLSQSLKRIADQFQKYL--------- | Jenson, J.M. *et al.* | none |
| M9 | --GRSQYEVIQELIRIGDIVLAYF--------- | Jenson, J.M. *et al.* | none |
| F10 | --GRRVVQIAAGLRRAGDQLEKYG--------- | Jenson, J.M. *et al.* | none |
| BAD | PNLWAAQRYGRELRRMSDEFVDSFKKGLPRPK- | BH3-only protein | none |
| X7 | --GQPLIWFGAQLRRGADEFAAQR--------- | Jenson, J.M. *et al.* | none |
| F4 | --GQRVVHIAAGLRRTGDQLEAYG--------- | Jenson, J.M. *et al.* | none |
| X1 | --GQTLIWYGASLRRYADEFAKQR--------- | Jenson, J.M. *et al.* | none |
| XF1 | --GRRVVWIGQGLKRLADEYHKYA--------- | Jenson, J.M. *et al.* | none |
| MX1 | --GRSQIWYVQELVRGGDVNHAYR--------- | Jenson, J.M. *et al.* | none |
| MX7 | --GRSEIWYDQELVRSGDVNAAYR--------- | Jenson, J.M. *et al.* | none |

**Table S1. BAK-binding signal for peptides that interact with anti-apoptotic BCL-2 family proteins tested using yeast-surface display.** Residues in the L-x(4)-D/E BH3 motif are highlighted are bolded.

Residues that align with positions of BIM that bind into hydrophobic pockets on BAK are in orange; residues in green are exceptions to the residue types typically found in BH3 motifs.

| PEPTIDE | SEQUENCE | BAK:DF2 dTERMen score | BAK:DF3 dTERMen score | BAK:DM2 dTERMen score | FINAL RANK |
|---|---|---|---|---|---|
| | 1 \| 2 \| 3 \| 4 \|5 | | | | |
| | efgabcdefgabcdefgabcdefga | | | | |
| **SNTG2** | NATHEEVVHLLRNAGDEVTITVEYL | 53 | 49 | 48 | 1 |
| **TXNDC11** | TRELQELARKLQELADASENLLTEN | 49 | 53 | 50 | 2 |
| **BNIP5** | DAIIQMIVELLKRVGDQWEEEQSLA | 53 | 51 | 54 | 3 |
| **TRIM58** | KSRLVQQSKALKELADELQERCQRP | 48 | 60 | 61 | 4 |
| **POFUT2** | TRRSMVFARHLREVGDEFRSRHLNS | 52 | 58 | 62 | 5 |
| **DDX4** | FSKREKLVEILRNIGDERTMVFVET | 54 | 63 | 54 | 6 |
| **PXT1** | EEIIHKLAMQLRHIGDNIDHRMVRE | 56 | 61 | 56 | 7 |
| **MINA** | TVATRRLSGFLRTLADRLEGTKELL | 57 | 60 | 60 | 7 |
| **CASP3** | SWFIQSLCAMLKQYADKLEFMHILT | 62 | 63 | 62 | 8 |
| **PCNA** | SGEFARICRDLSHIGDAVVISCAKD | 61 | 64 | 64 | 9 |
| **FOLH1** | PFDCRDYAVVLRKYADKIYSISMKH | 63 | 69 | 62 | 10 |
| **TRPM7** | FERVEQMCIQIKEVGDRVNYIKRSL | 62 | 72 | 62 | 11 |
| **TERT** | LRGSGAWGLLLRRVGDDVLVHLLAR | 63 | 68 | 63 | 11 |
| **PURB** | FKAWGKFGGAFCRYADEMKEIQERQ | 69 | 71 | 73 | 12 |
| **SLC19A1** | ACGDSVLARMLRELGDSLRRPQLRL | 71 | 71 | 74 | 13 |
| **MRPL41** | MGVLAAAARCLVRGADRMSKWTSKR | 67 | 74 | 76 | 14 |
| **NBEAL2** | AELRLFLAQRLRWLCDSCPASRATC | 69 | 74 | 76 | 15 |
| **MCF2L** | VDSIRPKCQELRHLCDQFSAEIARR | 69 | 74 | 78 | 16 |
| **MCF2L2** | ADAIRPRCVELRHLCDDFINGNKKK | 80 | 70 | 82 | 17 |
| **SPNS1** | ISSYMVLAPVFGYLGDRYNRKYLMC | 75 | 83 | 81 | 18 |
| **FOXJ2** | PAKKMTLSEIYRWICDNFPYYKNAG | 79 | 75 | 82 | 18 |
| **RTEL1** | RVCPYYLSRNLKQQADIIFMPYNYL | 89 | 81 | 88 | 19 |

**Table S2. Energy scores for BAK:peptide complexes calculated using dTERMen and ranked from lowest (best) to highest.** The indicated sequence was scored on each of three BAK:peptide complex structures using dTERMen and energies were rounded to the nearest integer; lower energies reflect greater predicted affinity. Ranks on the different templates were consolidated into a single overall rank. Some sequences had similar energy scores and were assigned the same rank number.

| PEPTIDE | SEQUENCE | UNIPROT ID | RESIDUES |
|---|---|---|---|
| BNIP5 | DAIIQMIVELLKRVGDQWEEEQSLA | P0C671 | 244-268 |
| TRIM58 | KSRLVQQSKALKELADELQERCQRP | Q8NG06 | 219-243 |
| PXT1 | EEIIHKLAMQLRHIGDNIDHRMVRE | Q8NFP0 | 76-100 |
| BIM | MRPEIWIAQELRRIGDEFNAYYARR | O43521 | 142-166 |
| PUMA | EQWAREIGAQLRRMADDLNAQYERR | Q9BXH1 | 131-155 |
| TRPM7 | FERVEQMCIQIKEVGDRVNYIKRSL | Q96QT4 | 1201-1225 |
| TERT | LRGSGAWGLLLRRVGDDVLVHLLAR | O14746 | 131-155 |
| SLC19A1 | ACGDSVLARMLRELGDSLRRPQLRL | P41440 | 245-269 |
| CASP3 | SWFIQSLCAMLKQYADKLEFMHILT | P42574 | 213-237 |
| SPNS1 | ISSYMVLAPVFGYLGDRYNRKYLMC | Q9H2V7 | 105-129 |
| NBEAL2Δ254 | AELRLFLAQRLRWLCDSCPASRATC | Q6ZNJ1 | |

**Table S3. Known and candidate BH3 motifs in human proteins.** Residues in the L-x(4)-D/E BH3 motif are highlighted in red.

| PEPTIDE | pLDDT SCORE OF BH3 REGION | STRUCTURE OF BH3 REGION | ACCESSIBILITY OF BH3 INTERACTION WITHIN PROTEIN |
|---|---|---|---|
| BNIP5 | 50-70 | alpha helical/ disordered | very accessible |
| TRIM58 | >90 | alpha helical | very accessible |
| PXT1 | >70 | alpha helical | very accessible |
| BIM | >70 | alpha helical | very accessible |
| PUMA | >70 | alpha helical | very accessible |
| TRPM7 | 70-90 | alpha helical | less accessible |
| TERT | >90 | alpha helical | less accessible |
| SLC19A1 | 50-90 | alpha helical/ disordered | less accessible |
| CASP3 | N.A. | alpha helical | inaccessible |
| SPNS1 | >70 | alpha helical | inaccessible |
| NBEAL2 | n.a. | n.a. | n.a. |

**Table S4. AlphaFold structure predictions and BH3 accessibility.** pLDDT scores indicate the confidence of structural prediction and are classified as follows: >90 - very high confidence, 90>pLDDT>70 - confident, 70>pLDDT>50 - low confidence, and pLDDT<50 - very low confidence. A pLDDT <50 is predicted as disordered (Tunyasuvunakool et al., 2021). We manually classified sequences into three groups: very accessible, less accessible, and inaccessible based on the predicted structure and the predicted alignment error. *n.a. indicates we are currently working on running NBEAL2 on Alphafold.

| PEPTIDE | SEQUENCE | FUNCTION |
|---|---|---|
| | 1 \| 2 \| 3 \| 4 \|5 | |
| | abcdefgabcdefgabcdefgabcdefgabcdef | |
| | h0 h1 h2 h3 h4 | |
| BIM-RT | ----DMRPEIRIAQELRRIGDEFNATYARR----- | Activator |
| Y-BID | YSESQEDIIRNIARHLAQVGDSMDRSIPPGLVNGL | Activator |
| PUMA M3DI | -----EQWAREIGAQLRRIADDLNAQYERRRQEEQ | Activator |
| dF8 | ------SLLEKLAEYLAQMGDEINKKYVK------ | Activator |
| dM4 | -----DKTLEEIARWLARLALEIDKEI-------- | Activator |
| dM2 | -----APYLEQVARTLRKIGEEINEALR------- | Activator |
| BNIP5 | -----DAIIQMIVELLKRVGDQWEEEQSLA----- | Activator |
| PXT1 | -----EEIIHKLAMQLRHIGDNIDHRMVRE----- | Activator |
| Y-NOXA | ---YPAELEVECATQLRRFGDKLNFRQKLL----- | Weak activator |
| Y-NOXA C2dI | ---YPAELEVEIATQLRRFGDKLNFRQKLL----- | Weak activator |
| PUMA | -----EQWAREIGAQLRRMADDLNAQYERRRQEEQ | Weak activator |
| BAD | ---PNLWAAQRYGRELRRMSDEFVDSFKKG----- | Non-activator |
| HRK | -LGLRSSAAQLTAARLKALGDELHQR--------- | Non-activator |
| dF2 | ------SYIDKIADL*I*RKVAEEINSKLE------- | Inhibitor |
| Y-dF7 | ------SLLEKLAEELAQLADELNKKFEK------ | Inhibitor |
| dF3 | -------LLEKLAEELRQLADELNKKFEK------ | Inhibitor |
| dF4 | ------SLLEKLAEYLRQMADEINKKYVK------ | Inhibitor |
| BK3 | -----KSPLERLAEILEKVA*K*EIEKELGP------ | Inhibitor |
| BIMh3PcRT | ----DMRPEIRIAQELRRXGDEFNATYARR----- | Inhibitor |

**Table S5. Alignment of peptide sequences tested for activation.** Residues are colored according to **Table S1**.

| Peptide | Sequence | N | MRE at 222 nm deg*cm$^{2}$*(10dmol*N)$^{-1}$ | Temperature (°C) at 222 nm | % helical content | Function |
|---|---|---|---|---|---|---|
| BIM-RT | DMRPEIRIAQELRRIGDEFNATYAR | 26 | -2353 | 24 | 5 | activator |
| Y-NOXA C2dl | YPAELEVEIATQLRRFGDKLNFRQKLL | 27 | -7280 | 24 | 17 | activator |
| Y-BID | YSESQEDIIRNIARHLAQVGDSMDRSIPPGLVNGL | 35 | -6945 | 25 | 17 | activator |
| Y-BAX BH3 | YPQDASTKKLSECLKRIGDELDSNMELQRMIA | 32 | -10862 | 24 | 26 | activator |
| dF8 | SLLEKLAEYLAQMGDEINKKYVK | 23 | -11667 | 25 | 27 | activator |
| dF2 | SYIDKIADLIRKVAEEINSKLE | 22 | -12240 | 25 | 28 | inhibitor |
| dM2 | APYLEQVARTLRKIGEEINEALR | 23 | -14975 | 23 | 34 | activator |
| dM4 | DKTLEEIARWLARLALEIDKEI | 22 | -15820 | 23 | 36 | activator |
| PUMA M3dl | EQWAREIGAQLRRIADDLNAQYERRRQEEQ | 32 | -20890 | 24 | 50 | activator |
| Y-BK3 | KSPLERLAEILEKVAKEIEKELGP | 25 | -28848 | 25 | 67 | inhibitor |
| Y-dF3 | SLLEKLAEELRQLADELNKKFEK | 23 | -35916 | 25 | 83 | inhibitor |

**Table S6. Percent helical content does not correlate with functional differences between inhibitors and activators**. 15 μM of peptide was used in each experiment. N indicates the number of residues in the sequence. MRE stands for mean residue ellipticity.

| Peptide | N-termini | Sequence | C-termini |
|---|---|---|---|
| Y-PXT1 | Ac | YEEIIHKLAMQLRHIGDNIDHRMVRED | NH2 |
| BNIP5 | Ac | DAIIQMIVELLKRVGDQWEEEQSLAS | NH2 |
| Y-TRIM58 | Ac | YKSRLVQQSKALKELADELQERCQRPA | NH2 |
| dM2 | Ac | APYLEQVARTLRKIGEEINEALR | NH2 |
| dM4 | Ac | DKTLEEIARWLARLALEIDKEI | NH2 |
| dF8 | Ac | SLLEKLAEYLAQMGDEINKKYVK | NH2 |
| BID-Y | Ac | EDIIRNIARHLAQVGDSMDRY | NH2 |
| BIM | Ac | MRPEIWIAQELRRIGDEFNA | NH2 |
| PUMA | Ac | EQWAREIGAQLRRMADDLNA | NH2 |
| PUMA2A | Ac | EQWAREIGAQARRMAADLNA | NH2 |

**Table S7. Peptide sequences used for BH3 profiling assay.**

| | DM2 | DF2 | DF3 |
|---|---|---|---|
| **Resolution range** | 38.07 - 1.3 (1.347 - 1.3) | 44.29 - 1.3 (1.347 - 1.3) | 40.35 - 1.99 (2.061 - 1.99) |
| **Space group** | C 1 2 1 | C 1 2 1 | P 1 21 1 |
| **Unit cell** | 102.13 41.04 47.02 90 93.644 90 | 94.13 41.08 56.39 90 122.001 90 | 48.29 65.26 111.57 90 102.117 90 |
| **Total reflections** | 320911 | 289866 | 160024 |
| **Unique reflections** | 46424 (3181) | 43727 (2462) | 43172 (4019) |
| **Multiplicity** | 6.91(100.88) | 6.63(117.73) | 3.7(39.81) |
| **Completeness (%)** | 96.66 (91.73) | 96.6 (73.5) | 92.3 (82.8) |
| **Mean I/sigma(I)** | 17.30(1.29) | | 6 |
| **Wilson B-factor** | 20.24 | 15.75 | 22.26 |
| **R-merge** | .04(1.71) | .03(0.58) | 0.15(0.71) |
| **R-meas** | .04(1.85) | 0.36(0.65) | 0.17(0.82) |
| **R-pim** | | | |
| **CC1/2** | 1(0.7) | 1(0.87) | 0.99(0.79) |
| **Reflections used in refinement** | 46366 (4360) | 43706 (3441) | 43044 (4010) |
| **Reflections used for R-free** | 2006 (188) | 2006 (159) | 1988 (178) |
| **R-work** | 0.1782 (0.3845) | 0.1615 (0.3179) | 0.2456 (0.3342) |
| **R-free** | 0.1982 (0.3849) | 0.1822 (0.3799) | 0.2758 (0.3808) |
| **Number of non-hydrogen atoms** | 1646 | 1692 | 6322 |
| **RMS(bonds)** | 0.02 | 0.02 | 0.004 |
| **RMS(angles)** | 1.6 | 1.57 | 0.62 |
| **Ramachandran favored (%)** | 100 | 98.37 | 99.45 |
| **Ramachandran allowed (%)** | 0 | 1.63 | 0.55 |
| **Ramachandran outliers (%)** | 0 | 0 | 0 |
| **Rotamer outliers (%)** | 1.26 | 0 | 0.65 |
| **Clashscore** | 3.3 | 1.32 | 4.42 |
| **Average B-factor** | 34.83 | 24.61 | 28.46 |
| **Number of TLS groups** | 8 | 0 | 0 |

**Table S8. X-ray data collection and refinement statistics.** Values in parentheses are for the highest-resolution shell.

| Peptide | Function | $k_{off}$ ($s^{-1}$) | stdev $k_{off}$ ($s^{-1}$) | $k_{on}$ ($M^{-1}s^{-1}$) | stdev $k_{on}$ ($s^{-1}$) | $K_d$ (nM) | stdev Kd (nM) |
|---|---|---|---|---|---|---|---|
| dF4 | inhibitor | 7.5E-02 | 6.8E-03 | 3.1E-04 | 2.9E-05 | 244 | 2.1E+01 |
| dF3 | inhibitor | 8.7E-02 | 1.2E-02 | 3.1E-04 | 5.0E-05 | 288 | 6.9E+01 |
| dF2 | inhibitor | 8.3E-02 | 2.2E-02 | 2.2E-04 | 4.3E-05 | 394 | 1.2E+02 |
| BK3 | inhibitor | 1.6E-01 | 2.4E-02 | 3.4E-04 | 8.5E-05 | 493 | 1.4E+02 |
| dF7 | inhibitor | 4.0E-01 | 1.4E-01 | 1.3E-04 | 5.3E-05 | 3294 | 5.5E+02 |
| BNIP5 | activator | 1.6E-01 | 2.6E-02 | 5.8E-04 | 2.4E-05 | 424 | 1.8E+02 |
| dF8 | activator | 2.0E-01 | 2.8E-02 | 1.1E-04 | 6.2E-05 | 3462 | 4.2E+03 |
| dM2 | activator | 1.1E+00 | 4.3E-01 | 6.5E-05 | 2.6E-05 | 17296 | 5.9E+03 |

**Table S9. Biolayer interferometry kinetics for BH3 peptides binding to BAK show a range of affinities that do not correlate with activation function.** Peptides were made as $His_6$-SUMO-BH3 fusions and immobilized on tips for binding to soluble BAK as described in the methods. Binding kinetics were too fast to fit for dM4 (activator) and PXT1 (activator) peptides.

| Peptide | Function | $K_d$ (nM) | Stdev $K_d$ (nM) |
|---|---|---|---|
| dF4 | inhibitor | 547 | 7.0+01 |
| dF3 | inhibitor | 594 | 1.7E+02 |
| dF2 | inhibitor | 155 | 8.2E+01 |
| BK3 | inhibitor | 151 | 3.1E+01 |
| dF7 | inhibitor | 1813 | 4.7E+02 |
| BNIP5 | activator | 411 | 1.7E+02 |
| dF8 | activator | 916 | 5.3+0.2 |
| dM2 | activator | 1480 | 5.9E+02 |
| dM4 | activator | 5307 | 2.0E+02 |
| PXT1 | activator | 10823 | 3.8E+03 |

**Table S10. Fluorescence anisotropy measurements.** Peptides tested were those listed in **Table S9.**

| Peptide | Function | BLI $K_d$ (nM) | FP $K_d$ (nM) | Classification |
|---|---|---|---|---|
| dF4 | inhibitor | 244 ± 2.1E+01 | 547 ± 7.0+01 | tight binder |
| dF3 | inhibitor | 288 ± 6.9E+01 | 594 ± 1.7E+02 | tight binder |
| dF2 | inhibitor | 394 ± 1.2E+02 | 155 ± 8.2E+01 | tight binder |
| BK3 | inhibitor | 493 ± 1.4E+02 | 151 ± 3.1E+01 | tight binder |
| dF7 | inhibitor | 3294 ± 5.5E+02 | 1813 ± 4.7E+02 | medium binder |
| BNIP5 | activator | 424 ± 1.8E+02 | 411 ± 1.7E+02 | tight binder |
| dF8 | activator | 3462 ± 4.2E+03 | 916 ± 5.3+0.2 | medium binder |
| dM2 | activator | 17296 ± 5.9E+03 | 1480 ± 5.9E+02 | medium binder |
| dM4 | activator | n.a. | 5307 ± 2.0E+02 | weak binder |
| PXT1 | activator | n.a. | 10823 ± 3.8E+03 | weak binder |

**Table S11. Affinities determined using bio-layer interferometry vs. fluorescence anisotroy give consistent classifications of peptide binders of BAK.** Peptides were grouped into tight (< 800 nM), medium (800 nM – 3 μM) , and weak (> 3 μM) binders for comparison to account for variability among methods. We were unable to measure affinities of weak binders dM4 and PXT1 by BLI. Fast kinetics for dM2 made it difficult to determine accurate $k_{off}$ and $k_{on}$ rate constants.
